# The shift from life in water to life on land advantaged planning in visually-guided behavior

**DOI:** 10.1101/585760

**Authors:** Ugurcan Mugan, Malcolm A. MacIver

## Abstract

Other than formerly land-based mammals such as whales and dolphins that have returned to an aquatic existence, it is uncontroversial that land animals have developed more elaborated cognitive abilities than aquatic animals. Yet there is no apparent *a-priori* reason for this to be the case. A key cognitive faculty is the ability to plan. Here we provide evidence that in a dynamic visually-guided behavior of crucial evolutionary importance, prey evading a predator, planning provides a significant advantage over habit-based action selection, but only on land. This advantage is dependent on the massive increase in visual range and spatial complexity that greeted the first vertebrates to view the world above the waterline 380 million years ago. Our results have implications for understanding the evolutionary basis of the limited ability of animals, including humans, to think ahead to meet slowly looming and distant threats, toward a neuroscience of sustainability.

## Introduction

The emergence of vertebrates on to land over 350 million years ago was preceded by a massive increase in visual range when their eyes moved to the top of the head to look over the water surface and tripled in size (*1*). The optical difference between inland waters—where transitional tetrapods are thought to have emerged—and air resulted in more than a 100-fold increase in visual range, from about a body length to hundreds of body lengths ahead (*1, 2*). Within the greatly enhanced range of Devonian aerial vision was rich structure provided by vegetation (*3*) and other terrestrial features, providing complex visual scenes to animals (**Supplementary Fig. 1A–B**). In this study we will test the hypothesis that for visually guided behaviors, the increase in visual range and observed environmental complexity that accompanied the onset of terrestriality advantaged the evolution of neural circuitry for deliberation over multiple futures (*4, 5*).

This study builds on investigations into the neural basis of action selection that suggest the existence of two competing, distinct, and largely parallel decision making systems: habit-based action selection, and plan-based action selection (*6, 7*). These two control paradigms have primarily been associated with the lateral striatum and its dopaminergic afferents (*8–10*) for habit, and the interaction between hippocampus and the prefrontal cortex (nidopallium caudolaterale in birds (*11*)) (*12–19*) for planning. The similarity between lamprey (jawless fish that preceeded mammals by 560 million years) and mammalian basal ganglia in the direct and indirect pathways of the dopamine expressing striatal projection neurons (*20–22*) suggests that this structure—and thus the habit-based action selection system it supports—evolved very early on in vertebrate evolution.

In mammals, planning has been related to the phenomenon of nonlocal spatial representations in hippocampal activity. Two quintessential examples of this phenomena are vicarious trial and error and sharp wave ripple events, where activity in the hippocampal place cells indicates a forward progression from the animal’s current location towards the goal location (*23–25*). These imagined action sequences are thought to arise from the interaction between the hippocampus and the prefrontal cortex. It is hypothesized that the prefrontal cortex sorts through context-relevant options to determine the best outcome as it engages the hippocampus (*4, 16–19, 26*). Birds have a hippocampus with the same developmental origin (*27*) and similar anatomy (*28*) as mammalian hippocampus, and there is accumulating behavioral evidence that it plays a similar role in planning for this group of animals through interaction with the avian homolog of prefrontal cortex (*11, 15*).

Despite the wealth of research into neural systems involved in habit- and plan-based action selection, the role that increased visual range and environmental complexity might have played in altering the relative advantage of these two decision making systems has not yet been explored. To address this gap, here we provide the results of a sequence of cognitive evo-devo (*29*) simulations aimed at assessing the relative advantage of habit- and plan-based action selection within aquatic and terrestrial environments during a goal-directed visually-guided behavior, predator evasion while approaching a distant refuge. This behavior was chosen due to its ethological relevance and evolutionary importance, but its logic makes it a member of a larger category of tasks—dynamic environments with shifting and uncertain rewards—which have previously been shown to engage planning in mammals (*6, 23*). We use formalizations of habit- and plan-based action selection from reinforcement learning theory (*30, 31*) to control discrete moves of a single prey and predator in environments with variable complexity.

## Results

In both habit-based and plan-based action selection, action choices are dependent on the current state of the animal. Within the context of prey pursuit by a predator, the current state is composed of the spatial location of the prey and the predator. The prey’s knowledge about these two variables allows it to associate a given outcome with an action, enabling it to predict long-term reward. Biologically, we can assume that the hippocampus (and its functional homologs) provide information about the prey’s location, while sensory information and memory allow it to infer the location of the predator, and connections to the prefrontal cortex (*32–34*) or its homolog in birds (*35*) provides an estimate of value for a given action sequence.

The habit solution to the problem of long-term reward prediction (often referred to as “model-free” (*6, 8, 31*)) simply assigns a value to an action or state based on prior experience. Thus, the action or state values are divorced from their outcome, which causes inflexible responses to changed circumstances that would require re-valuation. This inflexibility, such as to reward devaluation, mimics the behavioral characteristics of habit-based action selection (*9, 36, 37*). Conversely, the planning solution (often referred to as “model-based” (*6, 31*)) relies on an ‘action-outcome’ knowledge structure to generate action sequences by simulating futures states and their expected outcomes. While this method is computationally expensive, it also creates flexible responses to changing circumstances, such as those caused by the movement of a mobile threat or opportunity. This flexibility and re-valuation mimics the behavioral characteristics of plan-based action selection (*23, 25, 38*).

To study the effects of how much planning an animal can do on performance and behavioral complexity, as a function of visual range and within-range environmental complexity, two different predator-prey scenarios were simulated (Fig. 1A–B). For both of these computational experiments, the predator was designed as a reflex agent (aggressive pursuit based on current state with some randomness), with the addition of a belief distribution regarding the likely location of the prey when it was out of view. The prey was configured to have either habit-based action selection, or plan-based action selection with a preset number of states that it can forward simulate. We were constrained to simulate planning in only one of these two animals due to the high computational burden of simulating planning in both (*39*). First, we simulated mid-water aquatic conditions (Fig. 1A) where the prey’s visual range was varied in a simple (open) environment (the coral reef condition is also considered in a subsequent analysis). Second, we simulated terrestrial conditions (Fig. 1B) by adding randomly generated obstacles until a pre-determined level of clutter density was reached (see Methods). Unlike the aquatic condition, visual range was limited only by the presence of occlusions. If an occlusion existed along a line between the prey and the predator, then they could not see each other.

**Figure 1:**
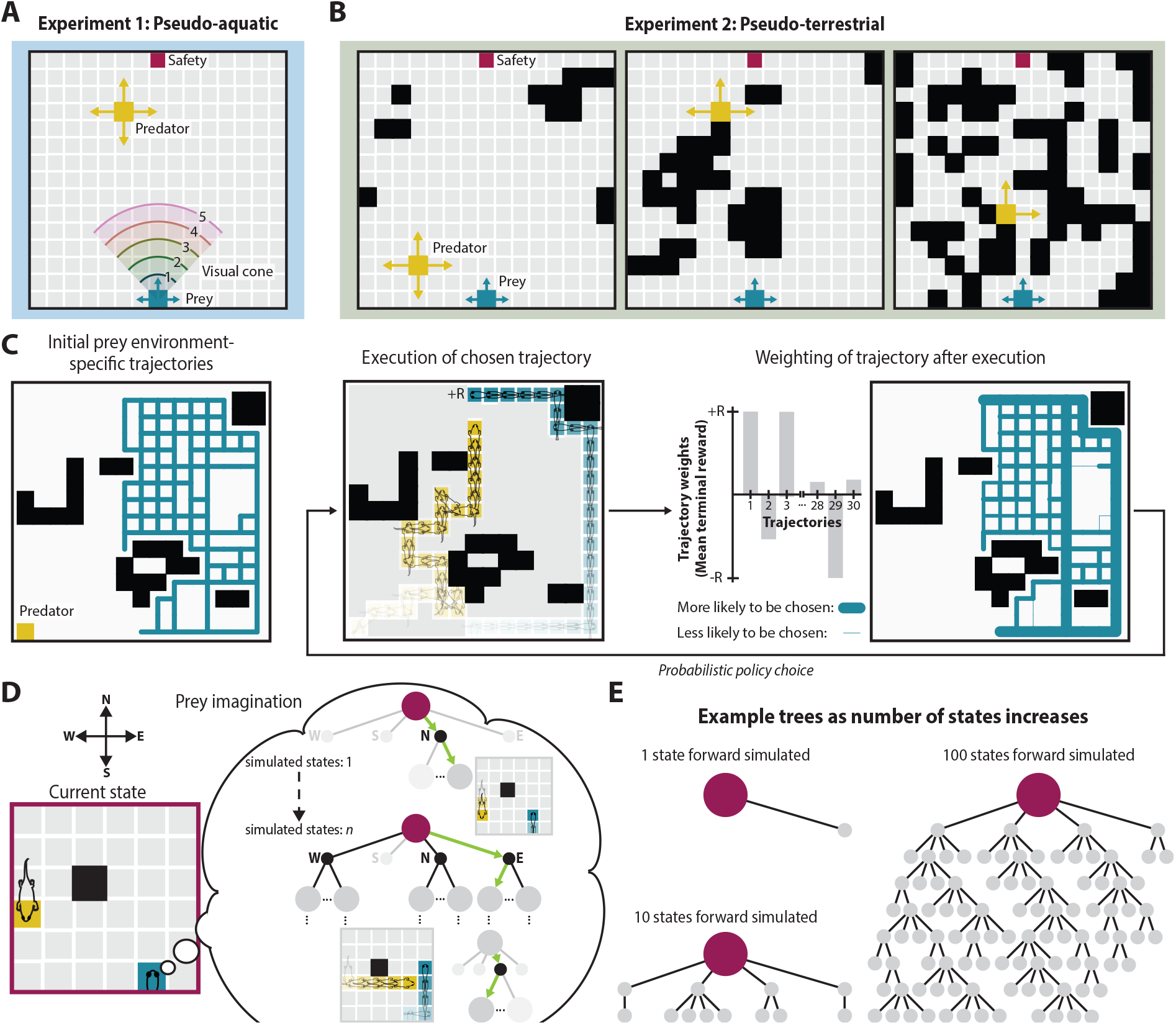
Environment models and algorithmic implementation of habit- and plan-based action selection. **(A–B)** Examples of the 2D environments used in the simulations; black squares represent obstacles. Examples of low, medium, and high clutter are shown for the pseudo-terrestrial experiment. **(C)** Schematic of habit-based action selection. Success paths for each environment (initially all weighted equally) are used as the initial condition for the illustrated loop in which one of a set of previously successful paths is randomly drawn with probability proportional to path weight. After executing the path, the weight of that path is adjusted to reflect the terminal outcome—higher weight for paths with higher terminal reward (see Methods). For illustrative purposes, we show the weight distribution after a number of trials. **(D)** Schematic of plan-based action selection, in which the prey imagines a tree of possibilities from the current state (dark red) by selecting virtual actions (selection shown in green, next state shown in dark gray, all other possible states shown in light gray). Example virtual action choices by the prey, reactionary virtual actions by the predator, and the resulting state is shown on the smaller grid. **(E)** Example trees grown given a specified number of states being forward simulated.

In both tasks, each trial had a fixed predator start location and number of states the prey could forward simulate, but both of these parameters were varied across trials. The prey’s start location and goal location (“Safety” in Fig. 1A-B) was fixed across all trials. Based on prior spatial navigation work with both fish and mammals (*40, 41*), we assumed the prey always knew its current spatial location. Both the predator and prey had a complete cognitive map of the space; the prey also had an accurate model of predator action selection.

### Habit-based and plan-based action selection

For this particular dynamic spatial navigation task, habit-based action selection exploits past action sequences that resulted in survival for a given visual range and/or environment (Fig. 1C). In real life, this would occur through trial and error over the lifetime of the animal or over evolutionary time; here, following past practice (*6*), we obtain these trajectories from plan-based action selection. Each action sequence within this set is weighted by the corresponding average execution outcome, thus causing the likelihood of selecting a specific action sequence to explicitly depend on whether it resulted in death or survival (*42*). While the probability of choosing an action sequence during habit-based action selection is increased if it resulted in past survival, the action sequence itself is not changed, which results in inflexible responses (*6*). Once an action sequence is chosen it is executed until termination (prey death or survival). In contrast, with plan-based action selection, within the imagination of the prey each virtual action is evaluated based on the virtual action’s possible outcomes. The prey thereby generates action sequences in imagination (Fig. 1D), which depends on the provided cognitive map and model of predator policy. This creates a tree structure (Fig. 1E) of virtual actions over the environment and their respective expected outcomes. This is somewhat similar to a chess player thinking through potential lines of play and counter-moves by an opponent, and possibly related to replay observed in spatial navigation tasks (*25, 38, 43*). After each action, the prey re-plans and thus re-valuates, which enables flexible responses to changes in predator location. The plan-based action selection method implemented in this study is based on a previously established efficient tree growth and search algorithm used within artificial intelligence approaches to games including AlphaZero (Monte-Carlo tree search (*44, 45*)).

### The utility of planning increases with visual range but cannot outperform habit in simple environments

In our pseudo-aquatic simulations of a predator-prey interactions (Fig. 1A), survival rate increases with the number of states forward simulated during planning, but the magnitude of this effect is proportional to visual range (Fig. 2A; Kruskal-Wallis test ***: *P <* 10^*−*7^). The utility of planning, as measured by the average change in survival rate across all tested planning levels, is significantly higher for long visual ranges (Fig. 2B; Mann-Whitney U (MWU) test with Bonferroni correction *: *P <* 0.03, Kruskal-Wallis test *P <* 10^*−*5^).

**Figure 2:**
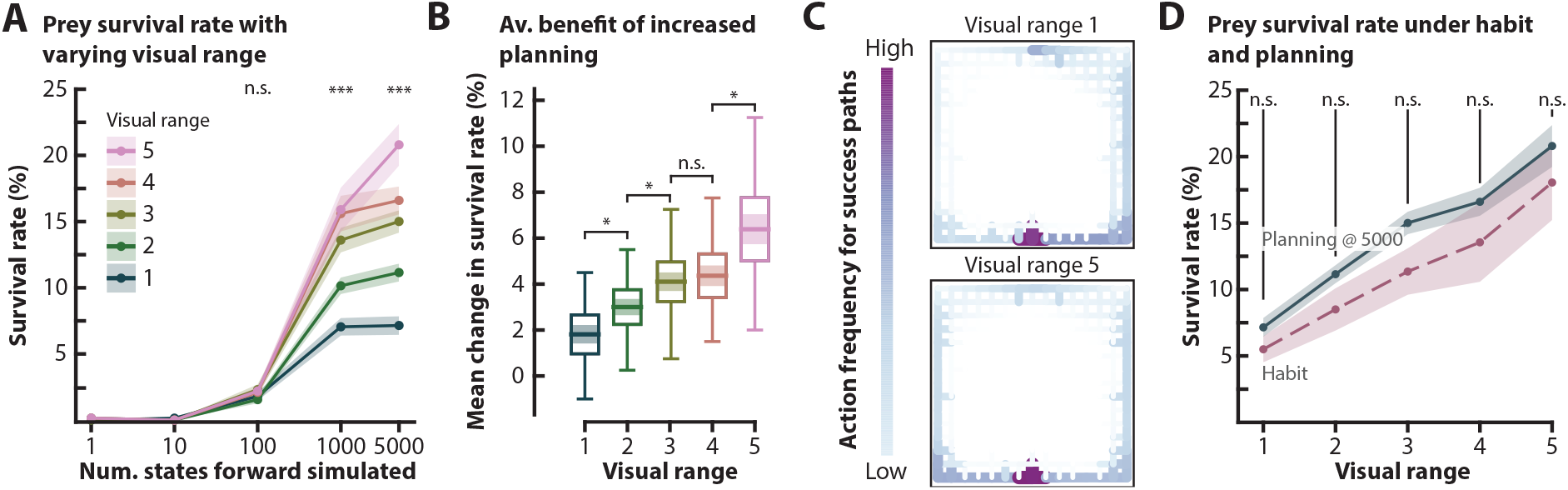
The utility of planning increases with visual range but cannot outperform habit in simple environments. **(A)** Prey survival rate in pseudo-aquatic environments (Fig. 1A) as a function of prey visual range and number of states the prey forward simulated across random initial predator locations (*n* = 20). The line plot shows the mean survival rate, and the surrounding fill indicates ± s.e.m. **(B)** Mean change in survival rate across all the planning levels shown in panel **(A)**—1, 10, 100, 1000, and 5000, calculated for each visual range. The horizontal line corresponds to the mean, the shaded region corresponds to the s.e.m., and the box corresponds to the 95% confidence interval of the mean. The lines extending from the box depicts the range of the data. **(C)** Heatmaps of all action sequences taken by the prey that resulted in prey survival at maximum planning level (5000 states forward simulated), with color density proportional to frequency. Color bar action frequencies range from 1−400, dependent on visual range (for trajectories across all visual ranges see **Supplementary Fig. 2**). **(D)** Mean ± s.e.m. of survival rate as a function of visual range at maximum planning (5000 states forward simulated). The planning data (teal solid line) is another representation of the plot shown in panel **(A)**. Mean survival rate as a function of visual range for prey that rely on habit-based action selection (pink dashed line). Fill indicates ± s.e.m. across tested predator start locations (*n* = 20). For all visual ranges there is no significant difference in survival rate between prey that use habit- and plan-based action selection.

In order to analyze the behaviors that result from increased planning across visual ranges, we quantified the frequency of actions between linked cells in those cases when the prey reached safety. The set of such action sequences are termed “success paths.” Interestingly, across all tested visual ranges, the simplicity of the environment results in stereotypical success paths (Fig. 2C). This suggests that high levels of planning provides the prey with a simple policy that takes the prey to the opposite wall from the estimated predator location (**Supplementary Movie 1**). Notably, these emergent successful action sequences resemble the wall-following behavior—or thigmotaxis— commonly observed in rodents in open-field tests (*46*). Habit-based action selection that uses these successful action sequences results in survival rates that are not statistically different from survival rates obtained with the maximum planning level (5000 states forward simulated) (Fig. 2D; One-way ANOVA all *P >* 0.05).

These results together strongly suggest that plan-based action selection, while advantaged in proportion to visual range, cannot improve performance over habit-based action selection in simple environments.

### Planning outperforms habit only in spatially complex environments

When the environment has very little clutter (entropy *<* 0.3), the predator speed and pursuit strategy (see Methods) restricts the prey’s survival rate (Fig. 3A), similar to what is observed in the pseudo-aquatic simulations. This survival rate restriction also limits the gain in survival with increases in the number of states the prey forward simulates before taking an action (Fig. 3B). Interestingly, at midrange levels of environmental clutter (entropy 0.4–0.6), both the prey survival rate (Fig. 3A), and the mean change in survival rate from increasing the number of states the prey forward simulates reaches its maximum (Fig. 3B; MWU test with Bonferroni correction Low-Mid: *P <* 10^*−*6^, Mid-High: *P <* 10^*−*5^). In environments with high levels of clutter (entropy *>* 0.6), both the effective size of the environment, and the number of possible escape routes to the safety position decreases, causing survival to be highly dependent on the predator’s initial location and the distribution of clutter. In both low and high entropy environments, there is no significant difference in the average change in survival rate across all planning levels (Fig. 3B; MWU test with Bonferroni correction Low-High: *P >* 0.05).

**Figure 3:**
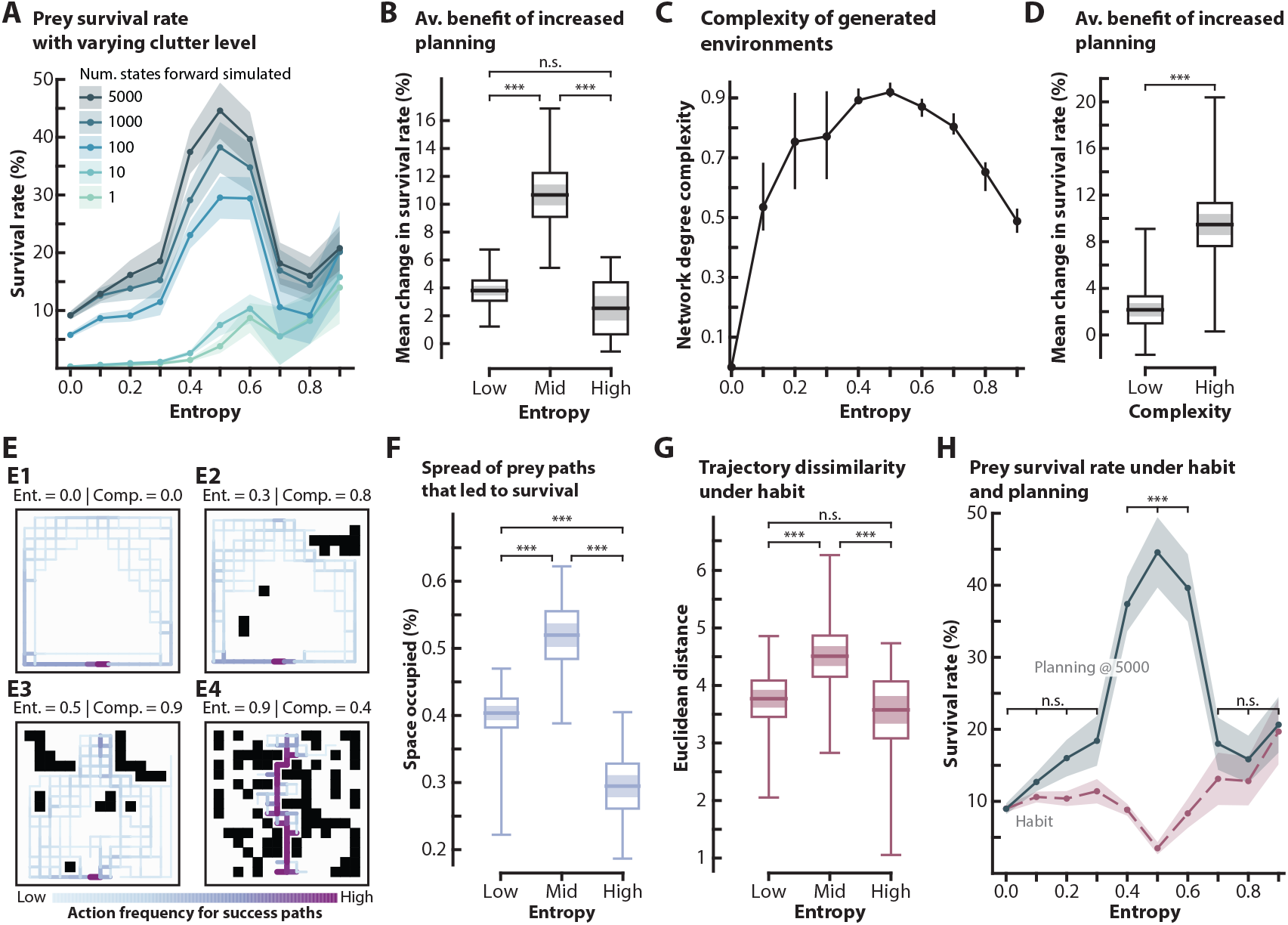
Planning outperforms habit only in spatially complex environments. **(A)** Mean survival rate versus clutter density (Fig. 1B) for a given number of states forward simulated. This is across different initial predator locations (*n* = 5), while fill indicates ± s.e.m. across randomly generated environments (*n* = 20). **(B)** Mean change in survival rate across all levels of forward states simulated shown in panel **(A)**—1, 10, 100, 1000, and 5000, calculated for each clutter level. Low: 0.0–0.3 entropy; Mid 0.4–0.6; High 0.7–0.9. The horizontal line corresponds to the mean, the shaded region corresponds to the s.e.m., and the box corresponds to the 95% confidence interval of the mean. The lines extending from the box depicts the range of the data. **(C)** Network complexity with respect to entropy. The line plot shows the mean complexity and the interquartile range (*n* = 20). **(D)** Mean change in survival rate across all the planning levels (see **(B)**), calculated for low complexity environments (entropy 0.0–0.3 and 0.7–0.9) and high (entropy 0.4–0.6). **(E1–E4)** Heatmaps of all action sequences taken by the prey that resulted in prey survival at the maximum planning level (5000 states), with color density proportional to frequency. Color bar action frequencies range from 1 for low, to 92 for the most saturated color, dependent on entropy level. Success paths in other environments are provided in **Supplementary Fig. 3**. **(F)** Path spread is defined as the percent of cells occupied by action sequences that resulted in prey survival, when the prey forward simulates 5000 states ahead (examples in **(E)**). High path spread indicates that the success paths are highly dissimilar. Plot representation as in **(B)**. Low, mid and high entropy ranges as in **(E)**. **(G)** Euclidean distance between action sequences that resulted in prey survival implemented by habit-based action selection. Greater distances imply higher dissimilarity between success paths. Plotted as in **(F)**. **(H)** Mean ± s.e.m of survival rate as a function of clutter level at maximum planning level (5000 states forward simulated). The planning data (teal solid line) is another representation of the plot shown in panel **(A)**. Mean survival rate as a function of environmental clutter for prey that rely on habit-based action selection (pink dashed line). Fill indicates ± s.e.m across all randomly generated environments (*n* = 20). For entropy levels 0.0–0.3 and 0.7–0.9 (low and high entropy groupings), there is no significant difference in survival rate between prey that use habit- and plan-based action selection.

The amount and distribution of clutter in the environment changes how visible each location is with respect to all other locations, causing different action sequences to be favored in relation to the predator’s movement. Since behavioral variability appears to be affected by the distribution of clutter via its affect on visibility at each cell, we used a network generated from these location visibilities. This visibility network connects all cells that do not have an occlusion along a connecting line of sight between cell centers (see Methods). The complexity of these networks, defined in terms of equivalence and diversity of cell visibilities, has two boundary conditions that are simple (*47*): fully connected environments (open), and unconnected environments (made up of only obstacles). However, between these edge cases, complexity (hereafter spatial complexity) increases until midrange levels of clutter, and then decreases for highly cluttered environments (Fig. 3C). It is in mid-entropy, high spatial complexity environments where planning becomes significantly more advantageous, indicated by the significantly higher average gain in survival rate from increased number of forward states simulated (Fig. 3D, MWU test: *P <* 10^*−*9^). In these environments, we observe high variability in the prey’s predator avoidance behavior (**Supplementary Movies 4–7**).

Similar to our analysis of behavior in pseudo-aquatic environments, we quantified behavioral variability in terms of action frequency, and created specific set of success paths for each environment (Fig. 2E). In accordance with our findings in aquatic environments (Fig. 2C), in low entropy terrestrial environments success paths under planning are stereotyped and thigmotactic (Fig. 3E1–E2, Fig. 3F, **Supplementary Movie 2**). In high entropy environments, characterized by high clutter and low spatial complexity, the amount of clutter constricts the profusion of success paths found at midrange entropy to one or two (Fig. 3E4, Fig. 3F, **Supplementary Movie 3**). Moreover, the prey’s survival strategy becomes less stereotyped as the number and spread of viable paths increases (Fig. 3E3, Fig. 3F; MWU test with Bonferroni correction Low-Mid: *P <* 10^*−*5^, Mid-High: *P <* 10^*−*10^). The spatial distribution of occlusions in these environments enables prey that use plan-based action selection to exhibit complex and flexible behaviors that strategically deploy occlusions to escape from the predator. Such behaviors can broadly be categorized as hiding and diversionary tactics, not unlike the broken-wing anti-predator tactic that is observed in birds (*48*) (**Supplementary Movies 4–7**).

The proportionality between the number of viable futures and environment spatial complexity affects the behavioral output and success of habit-based action selection. As expected, in low complexity environments, success paths under habit-based action selection are highly stereotyped (Fig. 3G, MWU test with Bonferroni correction Low-Mid: *P* = 0.009, Mid-High: *P* = 0.009), and not statistically different in Euclidean distance (MWU test with Bonferroni correction Low-High: *P* = 0.24). Notably, in both relatively open and highly cluttered low spatial complexity environments, there is no statistically significant difference in survival rate between habit- and plan-based action selection (5000 number of states forward simulated; Fig. 3H, One-way ANOVA Low: *P* = 0.15, High: *P* = 0.52). On the other hand, in complex environments, the high spread of success paths—the result of complex behaviors specific to predator choices—causes habit-based action selection to perform much worse than plan-based action selection, since habit-based action selection does not allow the rapid re-valuation of (Fig. 3H, One-way ANOVA: *P <* 10^*−*8^).

These results suggest that emergent strategies for survival are dependent on environmental properties interacting with dynamic contingencies—in this case, the position of the predator. Complex environments with dynamic contingencies—favoring visual tracking (Supplementary Text)—generate multiple viable futures. Our results predict significant selective benefit for animals with the circuitry to imagine these possibilities and discover their diverse values in comparison to animals with only habit-based action selection.

### Transitions in eigencentrality signal when there should be a switch between habit and planning

The low spread of success paths in simple environments suggests that the predator will follow a competing strategy that is similarly dependent on environmental properties. Complementary to our previous analysis, we quantified success paths for the predator by calculating the frequency of actions taken by the predator in episodes that resulted in predator success (capturing the prey) (Fig. 4A, **Supplementary Fig. 4**). The predator’s trajectories are similar to those observed in pursuit tasks in open environments with primates (*49*), and seem to arise as a result of easy access to predicted prey locations.

**Figure 4:**
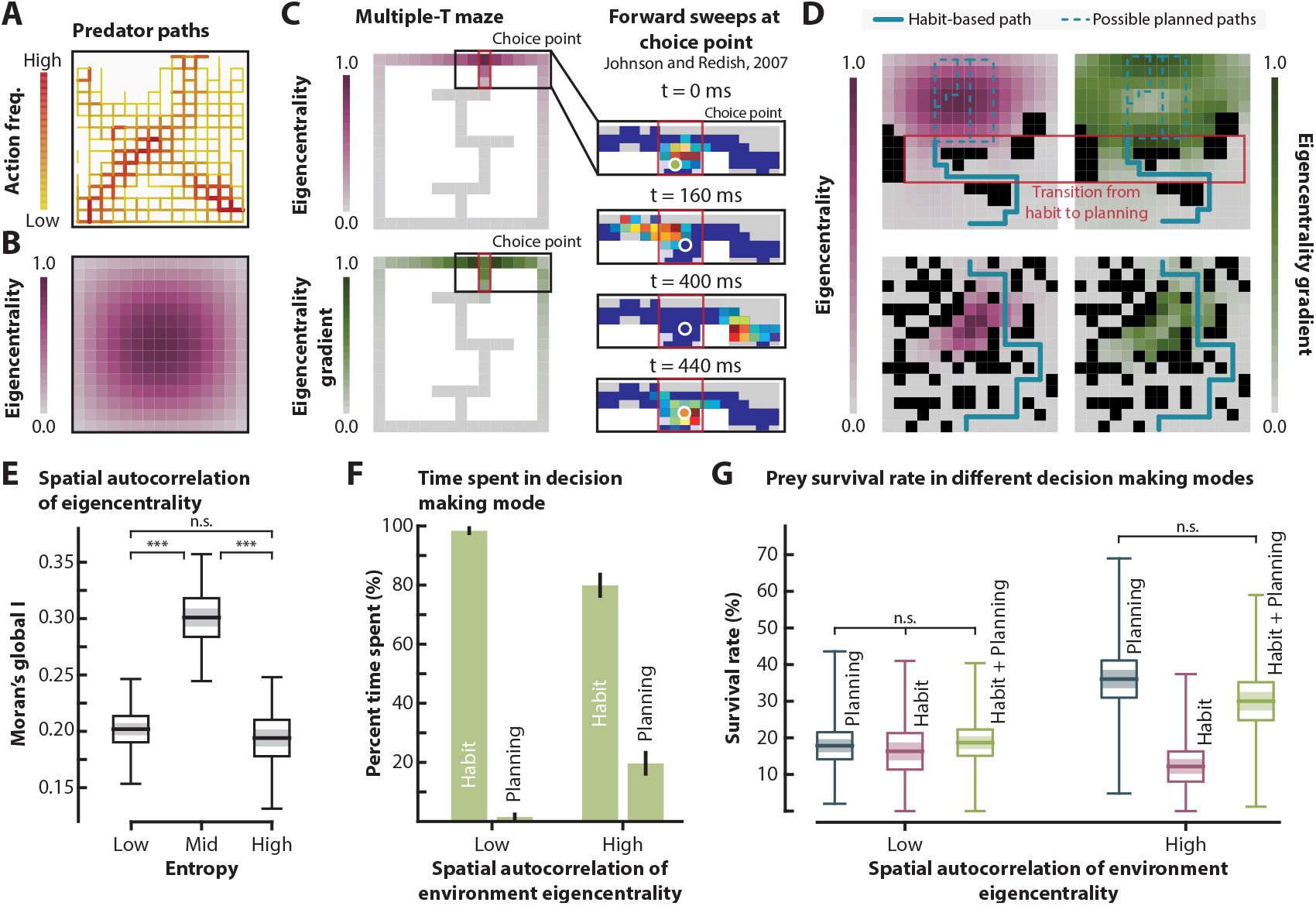
Transitions in eigencentrality signal when there should be a switch between habit and planning. **(A)** Heatmap of all predator success paths (capturing the prey before the prey reaches safety), with color density proportional to frequency. Color bar: low: 1, high: 90. **(B)** Eigencentrality of an open environment. Color density proportional to the eigencentrality of each cell. **(C)** Multiple-T maze overlaid with eigencentrality and eigencentrality gradient. Color density proportional to the eigencentrality and eigencentrality gradient of each cell of the quantized maze. Red box indicates the “choice” point where the rat pauses. Johnson and Redish (*23*) showed that the neural representation (reconstruction on the right) moved ahead of the animal while it paused at the choice point. The actual position of the animal is shown with a white circle. **(D)** Example environments (top row entropy = 0.5, bottom row entropy = 0.9) and their eigencentralities and eigencentrality gradients. Color densities of each cell are proportional to the metric. Transition regions from low to high eigencentrality based on change in gradient and value of eigencentrality are shown by the red box (for more examples see **Supplementary Fig. 5**). In our hybrid ‘planning+habit’ strategy, this transition region corresponds to a change in behavioral strategy from habit-based (solid teal line) to planning (dashed teal line). The prey success trajectories specific to these environments are shown in **Supplementary Fig. 6**. **(E)** Spatial autocorrelation (global Moran’s I) of the environment eigencentrality. Higher spatial autocorrelation indicates that the low and high values of environment eigencentrality are more spatially clustered. The horizontal lines correspond to the mean, the shaded regions correspond to the s.e.m, and the boxes correspond to the 95% confidence interval of the mean. The line extending from the box depicts the range of the data. **(F)** Average percent time spent in decision making regime (habit vs planning) when environments are grouped based on their spatial autocorrelation of eigencentrality. Low corresponds to the bottom 25%, which is largely made up of low and high entropy environments. High corresponds to the highest 75%, which is largely made up of mid-entropy environments. The error bars indicate ± s.e.m of percent time spent (*n*_low_ = 54, *n*_high_ = 50). **(G)** Survival rate for a prey that uses planning (blue), uses habitual control (pink), and uses hybrid control (green) based on environment eigencentrality (**(D)**; see Methods). Environment grouping is the same as **(F)**. Plot representation as in **(E)**.

For further analysis of this pattern, we quantified the connectedness of cells by eigenvector centrality (eigencentrality), which represents the weighted sum of direct connections (actions to and from a cell), as well as indirect connections of every length (*50*) (Fig. 4B) (see Methods). High eigencentrality implies that the cell is connected to cells, which themselves are highly connected (have high eigencentrality values). The few success paths in open (low entropy) environments employed by prey that explore 5000 future states, independent of prey visual range, are along cells of low eigencentrality (correlation between cell eigencentrality and action frequency in success paths: Spearman *ρ*_median_ = *−*0.54). Conversely, the predator success paths, independent of predator initial location, are more spatially distributed, and frequently occupy cells of high eigencentrality, which allows easy transitions to neighboring regions (Fig. 4A–B). Competitive sports analyses similarly point to the importance of reaching positions of high eigencentrality for rapid transitions to less connected regions (*51*).

In contrast, the emergence of complex behaviors in complex environments appears to be related to the distribution of eigencentralities. Unlike simple environments (Fig. 4B), which have a region of high eigencentrality that tapers away in all directions, spatially complex environments exhibit adjacent clusters of highly and poorly connected regions (correlation between spatial complexity and global Moran’s I: Pearson *ρ* = 0.61, *P <* 10^*−*21^) (Fig. 4D–E; MWU test with Bonferroni correction Low-Mid: *P <* 10^*−*6^, Mid-High: *P <* 10^*−*5^). In such environments simple action sequences—such as following cells with low eigencentralities—do not emerge (correlation between cell eigencentrality and action frequency in success paths: Spearman *ρ*_median_ = *−*0.05), resulting in poor performance for habit-based action selection.

We hypothesized that planning becomes more important, and results in higher behavioral variability, at regions of the space where there is a transition from low eigencentrality to high eigencentrality. These are found in environments that have adjacent clusters of low and high eigencentrality (Fig. 4D–E). Preliminary support for this hypothesis is found in the pattern of nonlocal hippocampal spatial representations that sweep in front of rodents at high-cost choice points (*23*), where there is a sharp change in eigencentrality (Fig. 4C). In order to examine this hypothesis further, we implemented a hybrid of habit- and plan-based action selection. Our hybrid strategy implemented habit-based action selection in low eigencentrality regions, which was switched to plan-based action selection when the prey moved into a region of high eigencentrality. In environments with low eigencentrality clustering (low and high entropy), behavioral control was rarely transferred over from habit-to plan-based (Fig. 4F). Consistent with our previous findings (Fig. 3H), in these environments all examined types of behavioral control strategies (habit-based, plan-based, and hybrid) performed similarly (Fig. 4G; One-way ANOVA *P* = 0.71). In contrast, in environments that have adjacent clusters of low and high eigencentrality (mid-entropy), plan-based action selection was engaged more often (Fig 4F; MWU test *P <* 10^*−*10^). While in these environments habit-based action selection performed much worse than plan-based (at 5000 states ahead), our hybrid strategy with switching based on environmental connectivity resulted in survival rates that were not significantly different from those attained with plan-based action selection (Fig 4G; MWU test with Bonferroni correction *P >* 0.05). Notably, the average decline in survival rate between a hybrid strategy with 75%– 85% of the time spent in habit-based action selection and the remainder in plan-based action selection (5000 states ahead) compared to 100% plan-based (5000 states ahead) was only 9%. The success of this hybrid strategy also agrees with prior research that indicates a need to transfer behavioral control strategy from habit-to plan-based action selection based on the agent’s uncertainty about state values (*6*). Within this context, the prey’s uncertainty increases with eigencentrality, due to the increase in regional openness and reachability.

### Natural terrestrial environments have high spatial complexity and advantage planning, while aquatic environments including coral reefs do not

In order to relate the randomly generated environments used here to natural environments, we used fractal dimension analysis (Fig. 5A), a commonly used approach to quantify the complexity of natural environments (*53, 58*). Intuitively, the fractal dimension (*D*) quantifies how the detail in a pattern (in this case a 2D environment) changes with the scale at which it is measured. Given that we are using 2-dimensional environments, the calculated fractal dimensions are between 0 and 2, where *D* = 0 corresponds to an empty, trivial environment, and *D* = 2 corresponds to environments with a large amount of fine structure. Inland and coastal aquatic environments have fractal dimensions in the range of 0.0–0.8 (see Methods), since the detail and complexity of whatever benthic structures or vegetation might be nearby cannot be seen due to high contrast attenuation (*1*).

**Figure 5:**
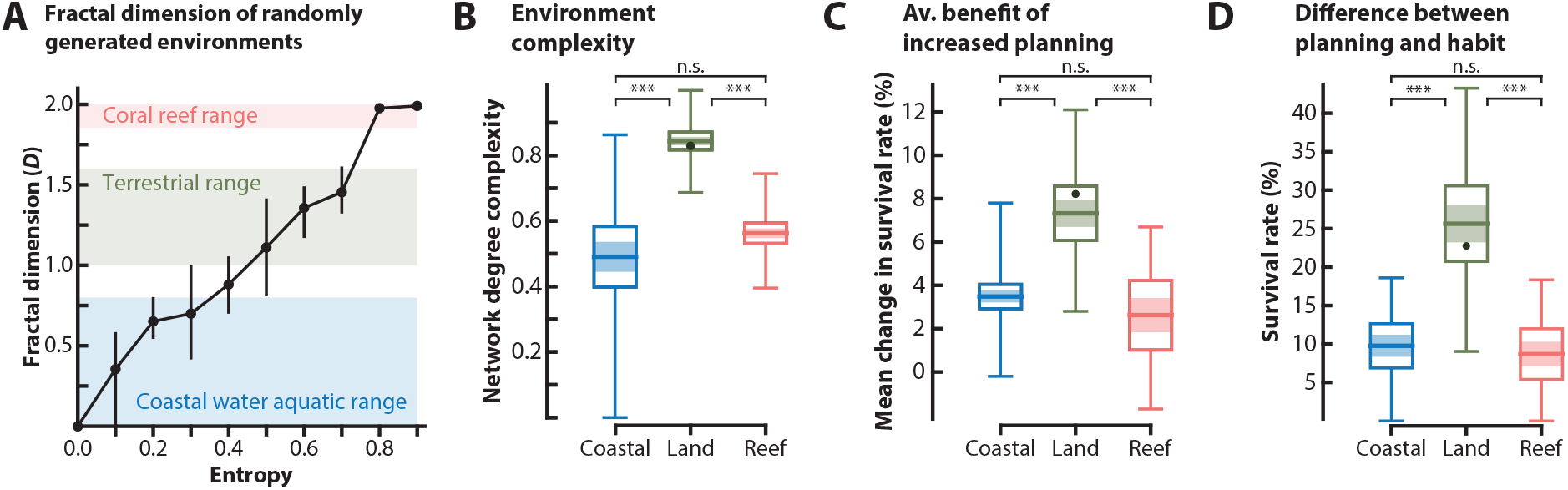
Natural terrestrial environments have high spatial complexity and advantage planning, while aquatic including coral reefs do not. **(A)** Distribution of fractal dimensions of generated environments. The line plot shows the mean fractal dimension and the interquartile range (*n* = 20). The blue fill shows the range of fractal dimensions (0–0.8, *n*_coastal_ = 66) that are observed in typical coastal aquatic environments (see Methods), the green fill shows the range of fractal dimensions (1.0–1.6, *n*_terrestrial_ = 66) that are observed in typical terrestrial environments (*52–54*), and the pink fill shows the range of fractal dimensions (1.9–2.0, *n*_coral_ = 39) that are observed in typical coral reefs (*55, 56*). **(B)** Environment network complexity (see Fig. 3C) of typical open water, terrestrial, coral reef environments. Dark green dot and band corresponds to mean environment complexity in environments that have a fractal dimension of ≈1.3 (peak human navigational performance (*57*)). The horizontal lines correspond to the mean, the shaded regions correspond to the s.e.m, and the boxes correspond to the 95% confidence interval of the mean. The line extending from the boxes depicts the range of the data. **(C)** The incremental benefit of planning (see Fig. 3B) of typical open water, terrestrial, and coral reef environments. Dark green dot and plot representation as in **(B)**. **(D)** The difference in survival rate between prey with high planning acuity, and prey that rely on habit-based action selection (see Fig. 3H) within typical open water, terrestrial, and coral reef environments. Dark green dot and plot representation as in **(C)**. In open water and coral reef environments there is no significant difference between habit- and plan-based action selection (One-way ANOVA: *P*_coastal_ = 0.1, *P*_coral_ = 0.25).

In contrast, structures and details can be viewed in coral reef environments, where the water is considerably clearer (see Methods), which results in high fractal dimensions (1.9–2.0) (*55, 56*). Similarly, the optics of terrestrial life enable the viewing of structural detail resulting in fractal dimensions between 1.0–1.6 (*52, 54*). We computed the fractal dimension of all our environments using a standard algorithm (*59*) (Methods) and used these ranges to categorize them into one of three major groups: coastal, terrestrial, and coral reef.

This categorization process shows that in terms of their fractal dimension, low entropy environments correspond to coastal aquatic environments, mid-entropy environments correspond to terrestrial environments, and high entropy environments correspond to coral reefs (Fig. 5B). This last grouping makes intuitive sense since coral reef habitats plus their surrounding water can be thought of as a high entropy space with multiple safety points (*60*) surrounded by a zero entropy open space. Based on this grouping, terrestrial environments appear to be significantly more spatially complex when compared to both coastal aquatic environments and coral reefs, which are statistically equivalent in spatial complexity (Fig. 5C; MWU test with Bonferroni correction Coastal-Terrestrial: *P <* 10^*−*7^, Terrestrial-Coral: *P <* 10^*−*13^, Coastal-Coral: *P* = 0.78). Given our earlier results, this indicates that terrestrial environments with dynamic contingencies advantage plan-based action selection, with advantage proportional to number of forward states simulated (Fig. 5D; MWU test with Bonferroni correction Coastal-Terrestrial: *P <* 10^*−*6^, Terrestrial-Coral: *P <* 10^*−*4^, Coastal-Coral: *P* = 0.52), and facilitates the emergence of complex behaviors. In contrast, coastal-like environments (fractal dimension corresponding to largely low entropy, 0.0–0.4) and coral reef-like environments (fractal dimension corresponding to high entropy, 0.8–0.9), advantage habit-based, stereotypical action sequences. There is unlikely to be significant advantage to planning in these variably cluttered (low and high) but spatially simple environments. A simple habit-based action selection that relies on a cognitive map, using previously successful action sequences, would likely perform just as well as high levels of planning (Fig. 5E; One-Way ANOVA Coastal habit-planning: *P* = 0.10, Coral habit-planning: *P* = 0.25, Terrestrial habit-planning: *P <* 10^*−*5^).

## Discussion

Our analysis shows that in simulated aquatic environments similar to those from which the early tetrapods emerged, or less turbid coral reef habitats, the simplicity of the environment generates stereotypical strategies. This allows habit-based action selection to perform as well as planning. Thus, there appears to be no selective benefit of planning circuitry to survive predator-prey interactions in these environments. This conclusion is buttressed by ethological analyses of predator-prey dynamics in open water or on open ground, which typically concern habitual and stereotyped responses. For example, analyses of aquatic prey behavior focus on escape trajectory analysis, where an escape trajectory is the initial segment of movement following detection of attack (*61–65*). Analyses of aquatic predator behavior typically focuses on strikes (*62, 66–68*). In aquatic contexts, the encounters are so short range that a specialized neuron called the Mauthner cell can be engaged to reduce the latency of movement by a few milliseconds, resulting in a higher chance of survival (*62*); this circuit disappears in the vertebrate line after amphibians (*61*). In most trials in simulated aquatic or low clutter terrestrial environments, the simulated prey demonstrate thigmotaxis, following the boundaries of the space (Fig. 2C, **Supplementary Movies 1–2**). Similarly, in studies of live animals in both aquatic and terrestrial contexts, retreating from exposed areas to edges after detecting increased predation risk is well documented (*69–72*), and is used as an assay of anxiety in laboratory experiments with fish and rodents (*46, 73*). Although the vast majority of the underwater predator-prey literature concerns reactive stereotyped behaviors, there are rare exceptions, such as cooperative hunting between moray eels and groupers (*74*), and triggerfish hunting behavior (review: (*75*)).

We found that in environments that have topographic features at a density similar to terrestrial environments, moment-to-moment changes in threat level due to changing positions of prey and predator relative to occlusions generates multiple viable futures. This advantages plan-based action selection, since only this mode of decision-making will rapidly propagate the expected value of taking an action. Our simulated encounters require both evasion from a predator and pursuit toward a distant goal, the refuge. Many of the interactions in the more complex midrange-entropy environments show the simulated prey using occlusions as a tool for hiding from the predator or luring them into a position from which they are better able to reach the refuge (**Supplementary Movie 5**, strategy 3). This strategic use of occlusions differentiates the relevent sensory ecology for this behavior, since only imaging systems like vision and echolocation will report occlusions, while auditory and olfactory systems require signal emitters, typically detected at much lower spatial and temporal resolution than useful in highly dynamic contexts (Supplementary Text). However, it should be noted that in our simulations, vision is only used to detect the predator, as both predator and prey are given an accurate cognitive map of occlusions and where the refuge is. Therefore, if a combination of non-visual modalities can provide sufficient localization and the animal has an accurate cognitive map, then an imaging modality may not be required for planning.

Within ethology, the use of occlusions to gain advantage during stalking is well known in Carnivora (*76–78*), though more common in felids than canids (*79*). Given the strategic use of occlusions by the simulated prey in our environments, we would expect similar behaviors to emerge had we simulated planning in the predator. The response of prey to predator pursuit in land animals is not as well resolved as the pursuit behavior of predators, being mostly described as fleeing behavior toward areas of cover (*69–71,80*) sometimes interspersed with freezing (*81*). While freezing was not an allowed action option for prey in our study, its functional role in concealment was observed to arise in our simulations in the form of the prey moving back and forth over a small area behind an occlusion between it and the predator (**Supplementary Movie 5**, strategy 3 & 5, **Supplementary Movie 6**, strategy 3, **Supplementary Movie 7**, predator location 3, strategy 3). A prediction for future investigation is that mammalian and avian prey may use occlusions as strategically as stalking predators from these groups.

We show that spatially complex environments have clusters of low and high eigencentrality. In these environments, we further show that using habit-based action selection within minimally connected space and switching to planning upon entrance to highly-connected regions performs as well as planning continuously. Vicarious trial and error, thought to reflect planning in rodents (*4*), has similarly been found to occur at transitions into highly-connected regions (Fig. 4C, (*23*)). Given the high computational burden of planning, a prediction for future investigation is that animals may switch their decision making mode accordingly. Prior research suggests that an increase in theta coherence between the hippocampus and prefrontal cortex is needed to sort through options as the hippocampus imagines potential outcomes (*4, 13, 26, 82, 83*). Interestingly, the coherence between the hippocampus and prefrontal cortex increases when rodents transition from a closed arm (low eigencentrality) to an open area (high eigencentrality) (*84*). This may portend a mechanism for switching decision making mode. It should be noted that eigenvector centrality has broad applicability to patterns of connectivity outside of our use of it as a measure of spatial connectivity (*85, 86*). As planning is similarly a concept that goes far beyond planning trajectories in space (e.g., (*16*)), it may be fruitful to examine whether transitions from habit-to plan-based action selection relate to eigenvector centrality changes in non-spatial contexts.

Given the strong evidence concerning early expansions in brain size of mammals related to olfactory processing (*29, 87*), it is possible that planning first evolved for situations that are not as continuously dynamic as visually-guided predator-prey interactions. Prior research has shown that there is a transition from plan-based to habit-based action selection after a novel set of contingencies arise but subsequently stabilize (*6, 88*). We will call such situations “habitizable,” while plan-based action selection for contingencies that do not stabilize—as considered in this work—are “non-habitizable.” An example of a habitizable use of planning would be an olfaction-dominant animal entering a new territory with relatively stable threats (e.g., the den of a predator) or opportunities (e.g., a large nest of insect prey). After update of the cognitive map, the planning system would be used initially for devising paths that avoid threats or result in faster access to the opportunities. Following some period of time these sequences would shift to habit-based action control.

Here, it is important to note a pattern found within the cognitive evo-devo (*29*) of mammals that is relevant to habitizable planning. That pattern is an inverse correlation between the size of limbic (LI) (olfactory bulb, olfactory cortices, amygdala, hippocampus, and septum) and isocortical (IS) components of the telencephalon (*29,89*). A somewhat similar pattern, remarkably, has been found across 62 species within the drosophilids, where an inverse resource allocation between vision and olfaction has been found (*90*). In both cases developmental constraints are suggested to play a role. In habitizable situations, the dependence on a cognitive map leads to an enlargement of LI components and planning can be cued in the near-field, such as by olfactory cues or low-acuity vision. However, given that logic, since cognitive maps are present in fish (*91, 92*) and reptiles (*93*), and they have near-field cues, they should have evolved planning in habitizable scenarios.

The evidence for planning in fish and reptiles thus far is at best sparse (*75, 91, 94, 95*). This could reflect insufficient work on testing its presence in these groups (*75, 94*), or that there is an added constraint that could make it difficult for them to evolve planning. In particular, plan-based action selection requires exponential computing time with the number of steps into the future being considered, while habit is constant time (Supplementary Text). Thus, in addition to needing significant sensory range to buy additional time before engagement with a mobile adversary—quite challenging in an aquatic context—the ability to plan may require a correspondingly larger amount of brain. The ability to perform plan-based action selection may therefore parallel what has been found for self-control, an ability that correlates with absolute brain size across 36 species of birds and mammals (*96*). Birds and mammals are endotherms, with brains that are ten times larger than those of ectotherms such as fish and reptiles at a body size of one kilogram, going up to forty times larger at large body sizes (*97, 98*). Furthermore, since all chemical reactions proceed faster with warming, the speed of movement and neural computation is generally higher in endotherms (Supplementary Text).

After the rise of habitizable planning in ancestral birds and mammals, with the subsequent evolution of visually-dominant birds with reduced olfactory bulbs (*99*), visually-dominant simians (*100, 101*), and terrestrial carnivores which hunt with long range vision and olfaction (*77–79*), we speculate that circuitry evolved to enable non-habitizable planning using rapidly changing distal cues interrogated by vision. In simians, this is correlated with a reduction in LI and an increase in IS (*29, 89*). The reduction in LI, whose overall volume is dominated by the hippocampus, could be related to an increase in distance between landmarks with long range vision, with more information extracted from a single gaze rather than through relational binding of numerous short-range landmark encounters. In terrestrial carnivores, a more balanced increase in both IS and LI occurs as they both use long range vision and olfaction during hunting (*29,89*). Our analysis suggests that strategic behavior in dynamic contexts through planning will play a larger role in these groups.

While this study is concerned with the utility of planning in situations where both predator and prey are moving rapidly, any dynamic context may benefit from planning, so long as the situation provides sufficient time such as through extended range or high acuity. For example, among invertebrates—where brain size is a less useful indicator (*102*)—the jumping spider has exceptional acuity that is 100 times better than the median for insects (*103*), similar to that of a pigeon. It shows evidence of planning while stalking its typically stationary prey, orb-weaver spiders (*104, 105*). One other invertebrate with exceptional visual acuity is the cephalopod, which are only lower than primates and raptors (*103*). They use neurally controlled pigment cells on their skin as if video displays, projecting detailed and rapidly changing patterns for deception during social interactions (*106, 107*). Here, a male will get between a female and rival, simultaneously displaying the female pattern to the rival and the male pattern to the female (*106*). This is a possible context for non-habitizable aquatic planning that does not need the extended sensory range required for planning in the predator-prey context, as it occurs at close quarters with slow changes in position.

A key question for future research into the mechanisms of planning is what constrains the temporal and spatial limits of the extended action sequences that animals are able to imagine prior to selection. Our results in computational cognitive evo-devo, examining how changes in sensory capacity and habitat in deep time affect the relative advantage of habit versus planning, can offer insights into cognition and the ability of animals—such as our own species—to make appropriate decisions for meeting distant and slowly looming threats at the margins of imaginative capacity.

## Acknowledgments

We thank Daniel Dombeck, Heydar Davoudi, Barbara Finlay, John Krakauer, and Lars Schmitz for helpful comments on an earlier draft.

## Author contributions

M.A.M. and U.M. designed research; U.M. and M.A.M. performed research; U.M. and M.A.M. analyzed data; M.A.M. and U.M. wrote the paper.

## Code and data availability

Code will be made available upon completion of the peer review process.

## Supplementary materials

Materials and Methods

Supplementary Text

Figs. S1 to S7

Tables S1 to S3

Captions for Movies S1 to S7

References

