## Supplementary Material for "The shift from life in water to life on land advantaged planning in visually-guided behavior"

**This PDF file includes**:

Materials and Methods

Supplementary Text

Figs. S1 to S7

Tables S1 and S3

Captions for Movies S1 to S7

References

### 1 Materials and Methods

This study concerns the performance of prey that only uses habit- or plan-based action selection. As such, for both pseudo-aquatic and -terrestrial simulations, we isolate the role of each control paradigm by assuming that the prey and the predator have previously learned the environment (where the boundaries are, where the occlusions are and the location of the safety). As the main acting agent, the prey is initialized with an environment and predator model, which allows the prey to forward simulate the actions of the predator. An episode is terminates if the prey reaches the safety location (survival), if the predator reaches the prey location (death), or if the number of steps exceeds the maximum number of steps allowed for an episode, here set to 200. For the pseudo-aquatic simulations, the number of steps before termination was 19 $\pm$ 6 (mean $\pm$ std). For pseudo-terrestrial simulations, the number of steps before termination across all clutter levels was 30 $\pm$ 17. The higher standard deviation arises because the number of steps varies greatly with the degree of clutter.

#### 1.1 Simulation 1: Pseudo-aquatic simulations

A virtual prey and predator act in an empty $15 \times 15$ virtual discretized environment (gridworld) (**Fig. 1A**). The aim of the prey, which starts at a fixed initial position at the middle of the bottom row of cells, is to get to the fixed safety position while being pursued by a predator. The prey receives reward of -1 at each step, -100 for dying and +1000 for surviving. The prey uses either habit- or plan-based action selection. Under plan-based action selection (see 1.1.1 **Simulation 1: Plan-based action selection and the evaluation of future states** for details) the prey has a predetermined number of states that it forward simulates before executing one action. As noted above, the forward simulation of states is dependent on an accurate environmental model (transitions between cells, and boundaries of the environment), and an accurate predator model (how the predator will move in space).

The predator is designed as a model-based reflex agent that selects actions based on the policy: aggressively pursue the prey with $75\%$ probability, and act randomly with $25\%$ probability. The predator is on average $1.5\times$ faster than the prey (moves 2 cells with $50\%$ probability), which falls within typical terrestrial predator speeds relative to prey of 1.2–2 (*1, 2*). The predator can observe the entire gridworld, and therefore knows the location of the prey at all time points. During aggressive pursuit, the predator chooses actions that minimizes the Euclidean distance between itself and the prey (if there is more than one action that meets this criteria, then an action from this set is chosen at random). During random action selection, if the prey is within the reach of the predator, the predator chooses actions that terminates the episode with predator success. Otherwise, the predator chooses a random action that keeps it within the confines of the gridworld. The predator spawns at a random initial position exclusive of the $3 \times 3$ region surrounding the prey, and the safety location. A total of $n = 20$ random predator locations were used for simulation 1 per visual range (1–5), and per number of states forward simulated (1, 10, 100, 1000, 5000). Survival rate was calculated over 100 episodes at a given predator spawn location, prey visual range, and prey planning acuity (for a total of 50,000 trials). For an overview of all general (planning or habit independent) parameters see **Supplementary Table 1**.

**Vision and partial observability.** The prey has a pre-specified visual cone that faces the direction of motion and extends outward 1–5 cells ahead (fixed at a single value per trial) (**Fig. 1A**). The prey always knows its own location within the gridworld, but only knows the predator location if the predator is inside of the prey's visual cone. If the predator is outside of the prey's visual cone, the prey samples a predator location from its belief state ($\mathcal{B}(., h_t)$) proportionate to the belief state distribution (**Supplementary Figure 7**) (for example, if the predator is believed to be at cell (8,12) with 90% probability, then 9 out of 10 draws from the distribution will be (8,12)). The number of samples (predator locations) the prey draws from its belief state is equal to the number of states the prey forward simulates. For each of these samples, the prey constructs a planning tree (**Fig. 1D–E**) and evaluates to the termination condition.

Upon spawning at the start location (7, 0) the prey rotates its visual cone to inspect its surrounding region. If the prey observes the predator during this sweep than it knows the predator initial location and therefore the current state. Otherwise, the prey's belief state is that the predator is located at all possible locations outside of its visual cone with equal probability. Until the first observation, the prey's belief state is propagated based on the prey's model of the predator movement; if the predator is then expected to be within the visual cone, but is not, then the set of belief states is correspondingly pruned and now the distribution reflects the new set of belief states (**Supplementary Figure 7**). In between two observations, the prey's belief state consists of all the possible places the predator might have moved given the location of the predator at the time of observation and the prey's model of the predator action-selection policy. For both the predator and the prey, staying at the current location was not an option; however, an approximation of holding station can occur through moving back and forth between two adjacent cells (e.g., Supplementary Movie 5, strategy 3).

One time step consists of: 1) The prey choosing an action with softmax decision maker (*3*) $a \in \mathcal{A}$ based on its forward evaluation (see 1.1.1 **CE1:Planning problem definition and evaluation of future states**). If that action brings the prey to the safety point then the episode is terminated (survival); 2) The predator choosing an action $a \in \mathcal{A}$ based on its pursuit policy (aggressive or random). If that action brings the predator to the prey position, then the episode is terminated (death); 3) The prey receiving an observation $o \in \mathcal{O}$ and reward $r = \mathcal{R}(s)$ from the environment; 4) The prey adding the action it chose and the corresponding observation to its history $h$; 5) The prey updating its belief state $\mathcal{B}(s, h)$ based on current history $h$; 6) The prey planning until a fixed planning acuity is reached.

##### 1.1.1 Plan-based action selection and the evaluation of future states

We formulate planning as a partially observable Markov decision process (POMDP) consisting of the following variables (*4*): a set of states $\mathcal{S}$ (prey and predator spatial location), a set of observations $\mathcal{O}$ (0 if the predator is not observed, cell number corresponding to predator location if the predator is observed), a set of actions $\mathcal{A}$ (cardinal directions: North, East, South, West), a list of action-observation pairs that constitutes the history $h$, a belief state $\mathcal{B}(s, h)$ specifying the probability distribution of the prey being in a state $s$ given history $h$, a reward function $\mathcal{R}(s)$ defined as the expected immediate reward for a given state $s$, and discount factor $\gamma = 0.95$ (*5*) that attenuates distal rewards.

The prey selects an action based on its policy $\pi(a|h)$, receives an observation, and collects rewards as it moves through states. Each state is associated with a value that denotes the expected total future reward starting from $s \in \mathcal{S}$ following a policy $\pi(a|h)$ ($\mathcal{V}^{\pi}(h)$). The maximum value that is achievable by any policy $\pi(a|h)$ gives the optimal value for that state ($\mathcal{V}^{*}(h)$). This value can be explicitly represented in terms of the reward function ($\mathcal{R}(s_t)$), and the belief state ($\mathcal{B}(s, h) = \mathbb{P}(s_t = s|h_t = h)$).

$$\mathcal{V}^{*}(h) = \max_{\pi} \left( \mathbb{E}_{\pi} \left[ \sum_{t=0}^{\infty} \gamma^{t} \mathcal{R}(s_t) \mathbb{P}(s_t = s|h_t = h) \right] \right)$$

Here, the prey's aim is to estimate $\mathcal{V}^{*}(h)$ by using its environmental model before taking an action (further discussed in 1.1.2 **Simulation 1: Algorithm of plan-based action selection**). The prey internally simulates its own actions, the reactionary actions of the predator, and the corresponding observations and rewards. These internal simulations are used to approximate the value function without explicitly calculating it.

##### 1.1.2 Algorithm of plan-based action selection

Forward simulation of future states is implemented by a tree-like planning system (Monte-Carlo tree search adapted for POMDPs (POMCP) (*4, 5*)), which relies on a previously learned model of the environment (boundaries and predator model). After an observation $o \in \mathcal{O}$ is received and a state sampled from the belief state $\mathcal{B}(., h_t)$, the prey begins planning from its current history $h_t$ to estimate the optimal value function $\mathcal{V}^*(h)$. Each node in the search tree, denoted by $T(h)$, has three elements associated with it: $\mathcal{B}(h)$ specifying a set of possible predator locations (which converges to a single state when the prey observes the predator), number of times a specific history $h$ has been visited ($N(h)$), and the expected value of an action and corresponding observation ($\mathcal{V}(h)$). $\mathcal{V}_{\text{init}}(h)$, and $N_{\text{init}}(h)$ are initialized to 0 for new nodes.

Planning tree construction and node value estimation in POMCP is divided into two stages: a tree-search policy that is on nodes with non-zero visit values (within-tree-search), and a rollout policy for nodes that have not previously been visited. After the evaluation of a state (predator and prey location), the node containing the first new history visited in the second stage is added to the search tree **Fig. 1E**. The planner uses partially observable UCT (PO-UCT) during the first stage within-tree-search, which selects actions based on Upper Confidence Bound (UCB1) (*6*); and a uniform random rollout policy during the second stage. PO-UCT has been proven to converge to the optimal value function (*5*),

$$\mathcal{V}(h) \rightarrow \mathcal{V}^*(h)\ \forall h \text{ as } N(h) \rightarrow \infty$$

which implies that when the planning acuity is high, an action that is selected by softmax based on the search tree is the optimal action to perform.

After an action $a_t$ is selected with a softmax decision maker (*3, 7*) and an observation $o_t$ is received from the environment, the planning agent's history is updated to reflect the new sample $\langle a_t, o_t \rangle$. The start node of the search tree and the associated belief state is updated to reflect the current history. The rest of the tree is pruned, since all other simulated histories are no longer representative of possible futures.

###### 1.1.3 Habit-based action selection

To model habit-based action selection (**Fig. 1C**), we implemented a variant of the PRQ-Learning algorithm (*8*). For each visual range, a policy library $L = \{\Pi_1, ...\Pi_n\}$ was created based on the prey success paths—prey going from the initial position to safety without being captured. These success paths were taken from policies implemented by the prey using plan-based action selection at 5000 states forward simulated. A policy $\Pi_k \in L$ was chosen by a softmax decision maker: $P(\Pi_k) = \frac{\exp^{\tau W_k}}{\sum_{p=0}^{n} \exp \tau W_p}$, where $W_k$ is the reuse gain of implementing the chosen policy, and $\tau$ is the temperature parameter. Initially all policies in the library are given zero weight. During policy implementation the prey does not deviate from the prescribed action sequence. After the implementation of the chosen policy $\Pi_k$, the reuse gain ($W_k$) is weighted by the total discounted reward $R$, and the number of times the policy $\Pi_k$ has been chosen ($N_k$): $W_k = \frac{W_k N_k + R}{N_k + 1}$. Initial values and parameters are provided in **Supplementary Table 3**. The predator action-selection policy was the same as the one implemented in the planning task (see 1.1 **Simulation 1**).

###### 1.1.4 Statistics

Significance among groups were tested by using one-way ANOVA and Mann-Whitney U test with Bonferroni correction where applicable. All significance indicators follow: n.s.: $P \geq 0.05$, *: $0.01 \leq P < 0.05$, **: $0.001 \leq P < 0.01$, ***: $P < 0.001$

In **Fig. 2B**, the incremental benefit of planning is defined as the average difference in survival rate between the tested 1, 10, 100, 1000, 5000 states forward simulated (e.g. difference in survival rate between 1000 and 100 states forward simulated) for a given visual range. Due to a non-uniform increase from planning acuity 1000 to planning acuity 5000 (difference is not 1 when converted to $\log$), a linear relationship was assumed, and the calculated difference was multiplied by 2.

#### 1.2 Simulation 2: Pseudo-terrestrial

A virtual prey and predator act in a $15 \times 15$ virtual discretized environment (gridworld) (**Fig. 1B**), that features randomly added clutter with controlled density. A total of $n = 20$ random environments, with $10$ levels of clutter—quantified by environmental entropy (see 1.2.1 **Simulation 2: Environment generation with randomized clutter** for details)—were generated. The predator and prey model used for this experiment are the same as 1.1 **Simulation 1**. The aim of the prey and the predator are kept the same as in 1.1 **Simulation 1**. The prey's plan-based action selection was based on the algorithm described in 1.1.1 **Simulation 1: Plan-based action selection and the evaluation of future states**.

The predator spawns at a random initial position exclusive of prey start location, safety position, and occlusions. A total of $n_{\text{pred}} = 5$ random predator locations were used for simulation per random environment ($n_{\text{env}} = 20$), per clutter level ($n_{\text{entropy}} = 10$: 0.0–0.9 in steps of 0.1), and per number forward states the prey was allowed to plan over ($n_{\text{forward simulation}} = 5$: 1, 10, 100, 1000, 5000). Survival rate was calculated over 50 episodes at a given predator spawn location, environment, clutter level and number of states forward simulated (for a total of 250,000 trials). A trial terminated if the prey reached the safety point, the predator moved the prey location, or if the episode reached cut-off point.

**Vision and partial observability.** Unlike 1.1 **Simulation 1**, the prey can see the entire environment except where blocked by occlusions (**Fig. 1B**). If an occlusion exists on the ray (Bresenham's line algorithm (*9*)) between the predator and the prey, the prey samples a state from its belief state ($\mathcal{B}(., h_t)$). Initially, if the predator is not observed by the prey (behind an occlusion), the prey's belief state is all possible locations that are unobservable from the given prey location. Until the first observation, the prey's belief state is propagated then pruned based on the prey's model of the predator movement and locations that are hidden from the prey's position. In between observations, the prey's belief state constitutes all the possible places the predator might have moved (based on the prey's model of predator movement) given the location of the predator at the time of observation and all the locations that are not visible from the prey's position.

The existence of occlusions impedes both the prey's and the predator's line of sight. Therefore, the predator knows the exact location of the prey if an occlusion is not present on the ray between the predator and the prey. The predator keeps track of the prey location while the prey is in view. When the prey is hidden, the predator propagates the prey's last known location randomly within the gridworld (exclusive of occlusions) to form a belief state. During aggressive pursuit, if the prey is within view the predator uses the actual prey location to choose an action that minimizes the Euclidean distance. If the prey is hidden, the predator aims to minimize its Euclidean distance to a randomly sampled prey location from the belief state. During random action selection, the predator chooses a random action that does not move the predator to an occlusion or outside of the gridworld.

The sequences of computations that occur in one time step are the same as 1.1 **Simulation 1**.

##### 1.2.1 Environment generation with randomized clutter

The entropy of a general $m \times n$ discretized environment can be calculated by treating the discretized environment as a binary matrix, where 1's represent occlusions, and 0's represent unoccupied cells. The entropy of such an environment ($\mathrm{Ent}(g)$) can be written as:

$$\mathrm{Ent}(g) = -\frac{\sum_{i=1}^{m}\sum_{j=1}^{n}\mathbb{I}(g_{i,j}=0)}{mn}\log\left(\frac{\sum_{i=1}^{m}\sum_{j=1}^{n}\mathbb{I}(g_{i,j}=0)}{mn}\right) - \frac{\sum_{i=1}^{m}\sum_{j=1}^{n}\mathbb{I}(g_{i,j}=1)}{mn}\log\left(\frac{\sum_{i=1}^{m}\sum_{j=1}^{n}\mathbb{I}(g_{i,j}=1)}{mn}\right)$$

where $g_{i,j}$ refers to the value at the $i^{\mathrm{th}}$ row and the $j^{\mathrm{th}}$ column.

In generating the occlusions for the environment we assume a random walk policy of a random length that starts at an unoccupied random position. The number of random walks performed can at least be 1 and at most be the number of occlusions for a given entropy, here denoted as $k$. The total number of random walk lengths $l$ must equal to $k$: $\sum_{1\leq l\leq k} l = k$. This process is repeated if a path from the fixed prey position to the fixed safety position does not exist (via the A$^*$ algorithm (*10*)).

##### 1.2.2 Environment complexity analysis

If an occlusion exists on the ray between the predator and the prey, both the prey and the predator are hidden from each other. By using the above principle we created a *visibility network* $G_v = (V_v, E_v)$ for all randomly generated environments (**Supplementary Figure**). Vertices in this visibility network represent individual cells in the gridworld. An edge $e_{i,j}$ exists between two vertices $\{v_i, v_j\}$ if an occlusion does not exist on the line between the two vertices, which is based on the same visibility ray used in determining the prey's and the predator's current observation. This can formally be written as:

$$E_v = \{\{v_i, v_j\}\,|v_i \in V_v, v_j \in V_v, v_i \neq v_j \text{ and } l(v_i, v_j) \cap O = \emptyset\}$$

where $l(v_i, v_j)$ determines the vertices that fall on the line between $v_i$ and $v_j$, and $O$ refers to the set of occlusions specific to the environment.

Each vertex $v_i$ has a degree $\deg(v_i)$ that specifies the number of vertices that are connected to the vertex $v_i$. With such a network transformation, an environment with $\mathrm{Ent}(g) = 0$ is a complete graph with vertex degree $\deg(v_i) = N - 1,\ \forall v_i \in V_v$. On the other hand, an environment that is only clutter is a disconnected graph with vertex degree $\deg(v_i) = 0,\ \forall v_i \in V_v$. Therefore, the complexity of a graph passes through a maximum and goes down to zero for complete and disconnected graphs (**Fig. 3**C). An argument in support of this complexity definition arises from Shannon's information theory applied to random graphs (*11*). Mathematically, the complexity of a network is defined as:

$$\alpha = \{deg(v_i) | v_i \in V_v\}$$
$$H(\alpha) = -\sum_{i=1}^{|\alpha|} \frac{\alpha_i}{|\alpha|} \log\left(\frac{\alpha_i}{|\alpha|}\right)$$

###### 1.2.3 Eigenvector centrality of environments

Environment quantization into a grid structure lends itself to a network representation based on how the system is connected together internally. If we again assume each cell is a vertex, in order to represent the environment dynamics, we can now define the edges in terms of actions. In such a network $G_w = (V_w, E_w)$, an edge $e_{i,j}$ between two vertices $v_i$, and $v_j$ exists if there is an action connecting the two vertices. Similar to before we can formally write this as:

$$E_w = \Big\{ \{v_i, v_j\} \, | v_i \in V_w, v_j \in V_w, v_i \neq v_j \text{ and } p(v_i) \xrightarrow{a} p(v_j) \ \forall a \in \mathcal{A} \Big\}$$

where $p(v)$ returns the cell for vertex $v$.

Eigenvector centrality (eigencentrality) depends both on the vertex degree $\deg(v_i)$ and neighboring vertex centralities. The centrality score $x$ of a vertex $v_i$ is defined as (*12*):

$$x_{v_i} = \frac{1}{\lambda} \sum_{j=1}^{n} A_{v_i, v_j} x_{v_j}$$

where $\lambda$ is the largest eigenvalue of the adjacency matrix $A_{v_i, v_j}$.

##### 1.2.4 Habit-based action selection

The prey's habit-based action selection was largely kept the same as the one described for pseudo-aquatic simulations (1.1.3 **Simulation 1: Habit-based action selection**). However, while the backbone of the algorithm (e.g. weighting and choice of policy from the policy library) was kept the same, the policy library was initialized differently. For each environment all the action sequences that led to prey survival during plan-based action selection were pooled together. If initially the prey was able to observe the predator, the policy library was pruned to reflect action sequences that were generated during plan-based action selection for that particular predator location. If the predator is not initially in view, the prey used the aggregate policy library after removal of all policies in which the predator was initially visible. For more details see 1.1.3 **Simulation 1: Habit-based action selection**.

##### 1.2.5 Transitioning between habit-based action selection and planning

We combined habit-based action selection and planning to model the consequences on survival rate for prey that switch between habit- (see 1.2.4 **Simulation 2: Habit-based action selection**) and plan-based action selection (see 1.1.2 **CE1: Planning algorithm**) based on the eigencentrality of the environment (**Fig. 4D**). We grouped environments based on their spatial autocorrelation of eigencentrality (SAE) (**Fig. 4E**), in which environments with SAE below the $25^{th}$ percentile were labeled as "low", and environments with SAE above the $75^{th}$ percentile were labeled as "high". We then performed a habit/planning switching protocol within these environments.

The prey switched from habit- to plan-based action selection (with 5000 number of states forward simulated) when transitioning from a low eigencentrality region to a high eigencentrality region. During habit-based action selection, the prey used its knowledge about the next action to determine if the new location had a higher eigencentrality. Conversely, the prey switched from plan- to habit-based action selection when transitioning from a high eigencentrality region to a low eigencentrality region. During plan-based action selection, the prey compared the eigencentrality and gradient of eigencentrality at its current location to all other possible locations (details provided below). The transition regions were identified based on the magnitude of the normalized eigencentrality ($X_E$) gradient:

$$|\nabla X_E| = \sqrt{\frac{\partial X_E}{\partial x}^2 + \frac{\partial X_E}{\partial y}^2}$$

During habit-based action selection, given the prey's knowledge about the predator policy, the prey retained a belief state. The belief state was set to be the current state when the prey observed the predator. If the prey was not able to observe the predator, the belief state was randomly sampled and propagated based the prey's model of the predator. If and when the prey switched over to plan-based action selection at a given transition point, the prey's belief state generated during habit-based action selection was used to initialize the belief state used by the planner (see 1.1.1 **Simulation 1: Plan-based action selection and the evaluation of future states** for more details). During plan-based action selection the belief state was set by the planner. If the prey switched to habit-based action selection, the belief state used by the planner was migrated over to the habit-based controller. This process was repeated for multiple switches.

At the start of each episode the prey's first action was chosen based on the policy selected by the habit control paradigm (see 1.2.4 **Simulation 2: Habit-based action selection** for more details). Given the nature of habit-based action selection, the prey's next position was calculated based on the next action prescribed by the selected policy. During plan-based action selection, the prey's next position was a set, comprised of all allowable locations (e.g. not walls and obstacles). The prey's previous location $(x_{t-1}, y_{t-1})$, current location $(x_t, y_t)$, and next location (for an environment of size $n \times m$ and set of obstacles $O$) $(\tilde{x}_{t+1}, \tilde{y}_{t+1}) =$

$\{(x_{t+1}, y_{t+1})| \, m > x_{t+1} \geq 0, n > y_{t+1} \geq 0, (x_{t+1}, y_{t+1}) \notin O\}$ were used to identify transition points. A transition was defined as:

$$|\nabla X_E(x_t, y_t)| > \max\{|\nabla X_E(x_{t-1}, y_{t-1})|, |\nabla X_E(\tilde{x}_{t+1}, \tilde{y}_{t+1})|\}$$

The prey's action selection algorithm was switched from habit- to plan-based if the prey was at a transition point, and the current eigencentrality value was smaller than the next eigencentrality value ($X_E(x_{t+1}, y_{t+1}) - X_E(x_t, y_t) > \epsilon$). Conversely, the prey's action selection algorithm was switched from plan- to habit-based if the prey was at a transition point, and the current eigencentrality value was greater than the maximum of the next possible eigencentrality values ($X_E(x_t, y_t) - \max X_E(\tilde{x}_{t+1}, \tilde{y}_{t+1}) > \epsilon$). $\epsilon$ was set to 0.001, for all other parameters see **Supplementary Information Table 2 and Table 3**.

##### 1.2.6 Environment fractal dimension analysis

The fractal dimension of the randomly generated environments (see 1.2.1 **Simulation 2: Environment generation with randomized clutter**) was calculated by using the box counting algorithm (*13*). Simply, boxes of decreasing sizes are inserted into the environment, and the number of cells that include occlusions are counted for each box. The fractal dimension of a gridworld that uses the box counting algorithm can mathematically be written as:

$$D = \frac{\log(N_r)}{\log(1/r)}$$

where $N_r$ is the number of boxes that cover the pattern and $r$ is the magnification, or the inverse of the box size. Therefore, the slope of the line when $\log N_r$ is plotted on the y-axis and $\log(1/r)$ is plotted on the x-axis equals the fractal dimension of the environment. A linear regression for this log-log plot was fitted to calculate the fractal dimension of each environment.

We used a similar approach on eight underwater images that ranged from murky to clear water to estimate the observed fractal dimension in aquatic environments. The colored aquatic images

($I$) of sizes $M \times M \times 3$ were converted to grayscale, and partitioned into a grid with box sizes $s \times s$, where $M/2 \geq s > 1$. Based on the differential box-counting approach (*14*), $N_r = \sum_{i,j} n_r(i,j)$, where $n_r(i,j) = \max I(i,j)_k - \min I(i,j)_k + 1$ and $k$ refers to the box number in the third dimension. $N_r$ is counted for different values of box sizes $s$, which determines the scale $r$. By using the above equation, we estimated $D$ from the least squares linear fit of $\log(N_r)$ against $\log(1/r)$. This analysis revealed that aquatic environments have fractal dimensions that range from 0–0.8.

##### 1.2.7 Statistics.

For all environment groupings, environments with entropies below the $25^{\text{th}}$ percentile were categorized as "low", and similarly environments with entropies above the $75^{\text{th}}$ percentile were categorized as "high". Mid-level entropy was classified as entropies between "low" and "high".

In **Fig. 3B** the incremental benefit of planning is defined as described above for 1.1.4 **Simulation 1: Statistics**.

For environment groupings in **Fig. 3D**, environments with spatial complexities below the $25^{\text{th}}$ percentile were categorized as "low", and similarly environments with spatial complexities above the $75^{\text{th}}$ percentile were categorized as "high".

In **Fig. 4E**, the spatial autocorrelation of the environment eigencentrality was calculated by using global Moran's I. The weight matrix was set to be the inverse of the vertex distances (min. number of actions to get from $v_i$ to $v_j$). This creates a weight matrix with greater values for vertices that are closer together. Global Moran's I evaluates whether a set of given values and their locations are clustered, dispersed, or random. For this statistic, the null hypothesis is that the spatial distribution of feature values (eigencentrality score of a vertex) is random.

For environment groupings in **Fig. 5C–E**, environments were grouped based on their fractal dimensions. Environments with fractal dimension below 0.8 were categorized as open water aquatic, environments with fractal dimension between 1.0–1.6 were categorized as terrestrial, and environments with fractal dimension between 1.9–2.0 were categorized as coral reef. Environments within the range 1.2–1.4 (not including the end points) were categorized as environments in which peak human navigation performance occurs (*15*).

Significance among groups were tested by using one-way ANOVA, Kruskal-Wallis, and Mann-Whitney U test with Bonferroni correction where applicable. All significance indicators follow: n.s.: $P \geq 0.05$, *: $0.01 \leq P < 0.05$, **: $0.001 \leq P < 0.01$, ***: $P < 0.001$

#### 1.3 Calculations of visual range for coral reef fish

In order to calculate the visual range for a typical coral reef fish we used a clear water model (*16*). This water model has an attenuation length of 1.17 m at 575 nm and a Secchi depth of 5.92 m. Given the position of the eyes we simulated visual range for horizontal viewing, using horizontal radiance ($\theta = 90°$ relative to nadir) with solar photon travel path angle $\phi = 180°$. This causes the diffuse attenuation coefficent ($K_d$) to be $0$ across all wavelengths and depths (for more detail see MacIver et al. supplementary materials: Aquatic Firing Threshold (*16*)). For a more realistic underwater range, we used the contrast threshold of goldfish (*Carassius auratus*) to account for object invisibility due to exponentially attenuated contrast (for more details see MacIver et al. supplementary materials: Contrast Threshold for Aquatic Vision (*16*))

We used a sample of 211 diurnal species of teleost reef fish in 43 families with a size range of 0.044–0.638 m across individual species (data obtained from Schmitz et al. (*17*)). Most species in this sample were reef inhabitants living in clear marine environments, with only a few entering murkier brackish and muddy coastal waters. For our simulations we did not differentiate between the two, and used a water model representing the clarity of the deepest oceanic water. Based on the data, pupil diameter increased with body mass. Therefore, we binned the body masses into 4 equal frequency bins and caluclated the average pupil diameter for each bin. This resulted in pupil diameters of 2.13 mm, 3.08 mm, 4.08 mm, and 5.17 mm. For each of these pupil diameters we calculated visual range using horizontal spectral radiance data for clear water at depths 5 m, 10 m, and 15 m. The calculated visual ranges at these depths for a 30 cm black disk "mock preator" (*18*) for fish with the above pupil diameters viewing the predator horizontally are: 5.02 m, 4.84 m, and 4.66 m for a fish with pupil diameter 2.13 mm; 5.04 m, 4.88 m, and 4.72 m for a fish with pupil diameter 3.08 mm; 5.05 m, 4.89 m, and 4.77 m for a fish with pupil diameter 4.08 mm; 5.06 m, 4.91 m, and 4.80 m for a fish with pupil diameter 5.17 mm. Given the size range of these fish, this causes the visual range to be $\approx$7–114 body lengths.

##### 1.4 Computing environment

The computational resources for this work were provided by the Quest high performance computing facility at Northwestern University which is jointly supported by the Office of the Provost, the Office for Research, and Northwestern University Information Technology. The cluster is composed of 244 nodes of Intel Haswell E5-2680 processors with 128 GB memory/node, 184 nodes of Intel Xeon E5-2680 processors with 128 GB memory/node, 72 nodes of Intel Xeon Gold 6132 processors with 96 GB memory/node. Approximate runtimes: **Simulation 1** 2,000 total compute hours (20 hours on 100 Quest nodes); **Simulation 2** 300,000 total compute hours (3,000 hours on 100 Quest nodes).

#### 2 Supplementary Text

##### 2.1 Fractal dimension analysis

Fractal dimension ($D$) concepts have been widely used to understand geographical complexity (*19–22*). These studies of natural scene fractal dimensions have approached the problem by representing the scene in terms of either a surface or as a contour. One important characteristic of fractal dimension is that going from a surface to a contour representation reduces the estimated fractal dimension by 1 ($D - 1$), and visa versa. These studies have calculated fractal dimensions based on a top-down view of the environment. While seemingly counter-intuitive, the relationship between contour (side view) and surface fractal dimensions points to the interchangeability, robustness, and perspective independence of the analysis.

In our current study we have been working with 2-dimensional environments, which would make fractal dimension be between $0 \leq D \leq 3$. However, given that the only stipulation posed on occlusions is that they have to be tall enough to disrupt the line of sight of both the prey and the predator causes us to not have height, and thus surface information. In calculating the fractal dimension of our randomly generated environments, we treated occlusions as points on a contour line.

It is important to note that within our study, while occlusions blocked the line of sight, neither the prey nor the predator relied on vision to locate the occlusions. The acting agent's having a perfect map of the environment means that they were navigating and (for the prey) imagining trajectories in the environment from a perspective-independent representation. For the fractal dimension calculation (see Methods) we translate this into a top-down perspective. This is similar to a human navigation study in which the fractal dimension of the environment was calculated based on a top-down perspective of the environment to be navigated (*15*).

#### 2.2 The sensory ecology of planning

The specific regime of our study is non-habitizable planning—environments so dynamic that plan-based action selection, with its unique ability to rapidly propagate the value of future actions back to inform what the next action should be, is favored over habit-based action selection. In our study, the key factor advantaging plan-based action selection is the presence of a mid-range level of occlusions. Such occlusions, such as terrestrial features like hillocks (*23*), do not typically actively emit signals that can be strategically exploited, but rather are signal reflectors. This makes their use contingent on one of two possibilities: 1) the possession of an imaging modality—vision or echolocation; 2) the possession of an accurate and detailed cognitive map in combination with a non-imaging modality (or several in combination) that provides sufficiently precise localization of a moving threat or opportunity. Regarding the first possibility, vision is clearly used by many animals for this purpose (see Discussion). Echolocation, a more recent innovation in mammals (*24*), provides good resolution and fast update rates, but is range-limited by power requirements (*25*). Nonetheless, sound propagation through water is as favorable as light propagation through air, so underwater echolocators can gain substantial range advantage over aquatic vision (*25*). We would expect our results to generalize to aquatic and terrestrial mammals engaged in echolocation-guided behaviors provided range is sufficient. Regarding the second possibility, our study provisioned an accurate cognitive map to both predator and prey (Methods), so vision was only needed for predator localization—animals which emit olfactory and auditory cues. Given the olfactory dominance of many mammals including rodents and elephants (*26, 27*) with evidence of cognitive maps (*28, 29*), the use of a cognitive map in combination with a non-imaging modality seems quite likely to occur for survival in non-habitizable contexts. In the absence of a cognitive map, or given an incomplete or inaccurate cognitive map, than an animal may need to rely on habit-based action selection or on imaging modalities for planning.

In habitizable planning, where planning generates action sequences initially used for advantage over relatively stable features prior to being shifted over to the habit system, olfaction, audition, path integration, and somatosense are all relevant. Hearing provides range and fast updates, but spatial resolution of sound emitters is generally poor outside of owls that hunt by precise sound localization. Olfaction provides long range, a slow update rate, and variable resolution depending on whether the odor is on a substrate or dispersed in air. It may be immensely important for the generation and indexing of a cognitive map, particularly in many olfaction-dominant mammals such as rodents and elephants (*26–30*). Cognitive maps have many uses that are independent of planning, such as present location on a map (*31*) and corrections to a dead reckoning system (*32*). This may result in broader phylogenetic distribution of this trait than planning, with evidence for its presence in fish (*33, 34*), in turtles (*35*), and in insects (*36*).

#### 2.3 Ectothermy, reptiles, and the computational complexity of planning

The cognitive ability of reptiles is poorly understood, and there is little evidence of planning (*37, 38*). Yet, these animals have existed in the same complex terrestrial habitats, with aerial vision, as mammals and birds. We consider two possibilities to resolve this conflict, although clearly this does not exhaust all possibilities. The first is that these animals are able to plan and this is simply not known due to lack of study; the second is that the selective benefit of planning, while applicable to these animals, is not high enough to overcome an additional constraint.

There is certainly not enough evidence to settle this issue presently, but there are a number of considerations in favor of the "additional constraint" hypothesis related to the computational cost of planning. While computers process information differently from neural circuits, the differences in the algorithmic complexity of habit versus planning as it is formalized by reinforcement learning theory is instructive. With habit-based action selection, policy lookup time is constant and independent of the number of actions within the action sequence. Conversely, with plan-based action selection, the time needed for an action choice scales exponentially with the number of steps forward simulated.

Both mammals and birds are endotherms. In mammals, endothermy appears to have come about due to selection for increased aerobic capacity in Permian theriodont therapsids, many of which were active predators, resulting in extended capacity for prey pursuit and predator avoidance (*39*). A 10°C increase in temperature of muscle above ambient doubles the rate at which muscle can reach maximum power (*40*). Endothermy likely played an important role in greatly increasing the size and computational power of the brain (*41, 42*), and in increasing the sensitivity and temporal resolution of vision (*43*).

The flicker fusion frequency of the retina of swordfish, a highly active underwater predator that rapidly transitions between warmer surface waters and cold deep waters, rises from 5 Hz in 10°C water to over 40 Hz at 20°C (*43*). This is one explanation for why swordfish is one of only around 30 out of 30,000 species of bony fish that has a heating mechanism, in this case for its eyes and brain only. However, endothermy is rare even among active predators underwater, likely because heat is dissipated 3,000 times faster within water than in air. In addition to this barrier to endothermy underwater, any increase in aerobic capacity of animals that gill rather than breathe has to contend with water having only 1/30th the oxygen of air, while being 800 times denser. Gill ventilation therefore requires 800 x 30 = 24,000 times more mass flux assuming identical extraction efficiencies; even with the doubled efficiency of gills the mass flux is four orders of magnitude higher for respiration with water compared to air (*16, 44, 45*). Given the disadvantageous energy load due to rapid cooling and the difficulty of recouping this energetic cost through additional oxygen expenditure, for most underwater animals the heightened velocity of movement, brain power, and sensory performance of endothermy is out of reach. Yet these factors seem to be important for realizing the selective benefit of planning, which also suggests that planning is either unlikely to occur in underwater animals or occur in diminished form relative to birds and mammals.

### 3 Supplementary Figures

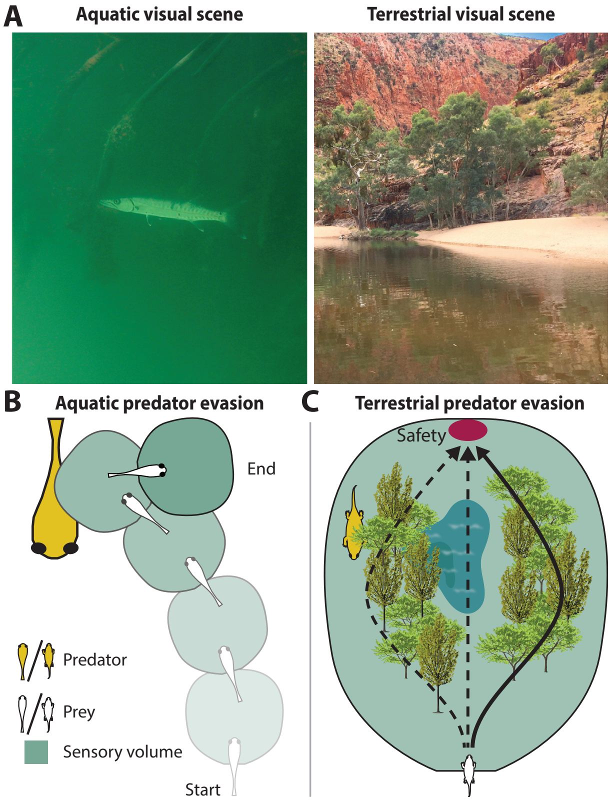


Figure 1: **Aquatic versus aerial visual scenes, and the hypothesis under test of how these habitats affect the utility of habit- versus plan-based action selection in a critical visually-guided behavior. (A)** Examples of aquatic and terrestrial visual scenes (from (*46*)). **(B)** The impact of sensory ecology on affording habit-based versus plan-based action selection. Because water rapidly absorbs and disperses light, aquatic behavior is typically short range and predator-prey interactions are at close quarters, requiring rapid and simple responses. **(C)** Terrestrial visually guided behavior can occur at long range due to the $\approx$100x increase in sensing range. This increase affords time for plan-based action selection wherein multiple possible futures are imagined (solid and dashed black arrows), followed by selection of the option with higher expected value (solid black arrow).

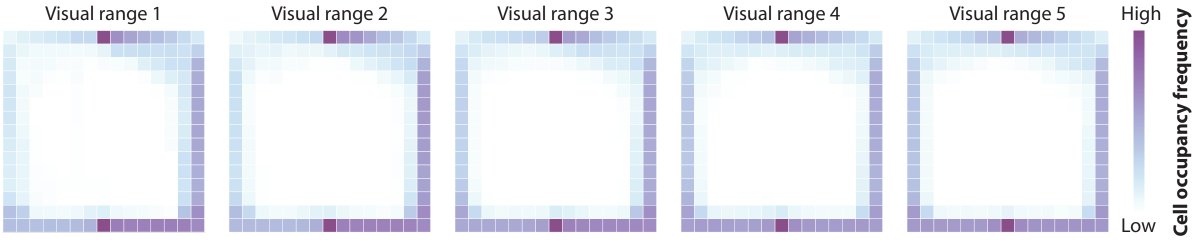


Figure 2: **Prey success trajectories with respect to visual range.** Heatmaps of all action sequences taken by the prey that resulted in prey survival at maximum planning level (5000 states forward simulated), with color density proportional to frequency. Color bar action frequencies range from 1–400, dependent on visual range. This representation shows that across visual ranges, the prey strategy remains the same. Longer visual ranges enable the prey to turn away from a sensed predator at a greater distance, allowing it to escape.

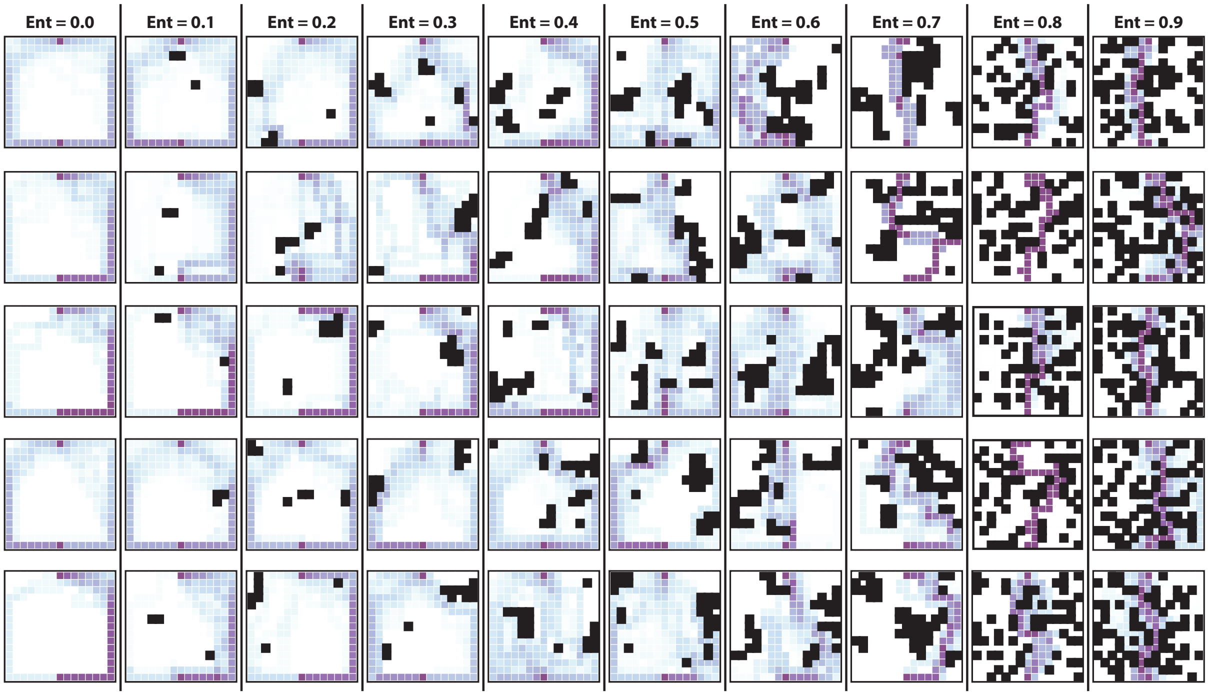


Figure 3: **Prey success trajectories with respect to environment entropy.** Heatmaps of all action sequences taken by the prey that resulted in prey survival at maximum planning level (5000 states forward simulated), with color density proportional to frequency. Color bar action frequencies range from 1–100, dependent on environment entropy. This representation shows that at low and high entropy success paths are fairly stereotypical. On the other hand, at midentropy environments, with the emergence of multiple viable futures, success paths are dissimilar.

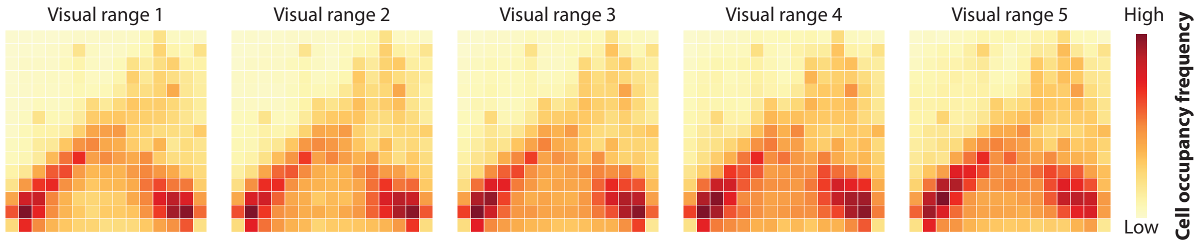


Figure 4: **Predator success trajectories with respect to prey visual range.** Heatmaps of all action sequences taken by the predator that resulted in predator capture of the prey, with color density proportional to frequency. Color bar action frequencies range from 1–500, dependent on visual range. This representation shows that across visual ranges, the predator strategy remains the same.

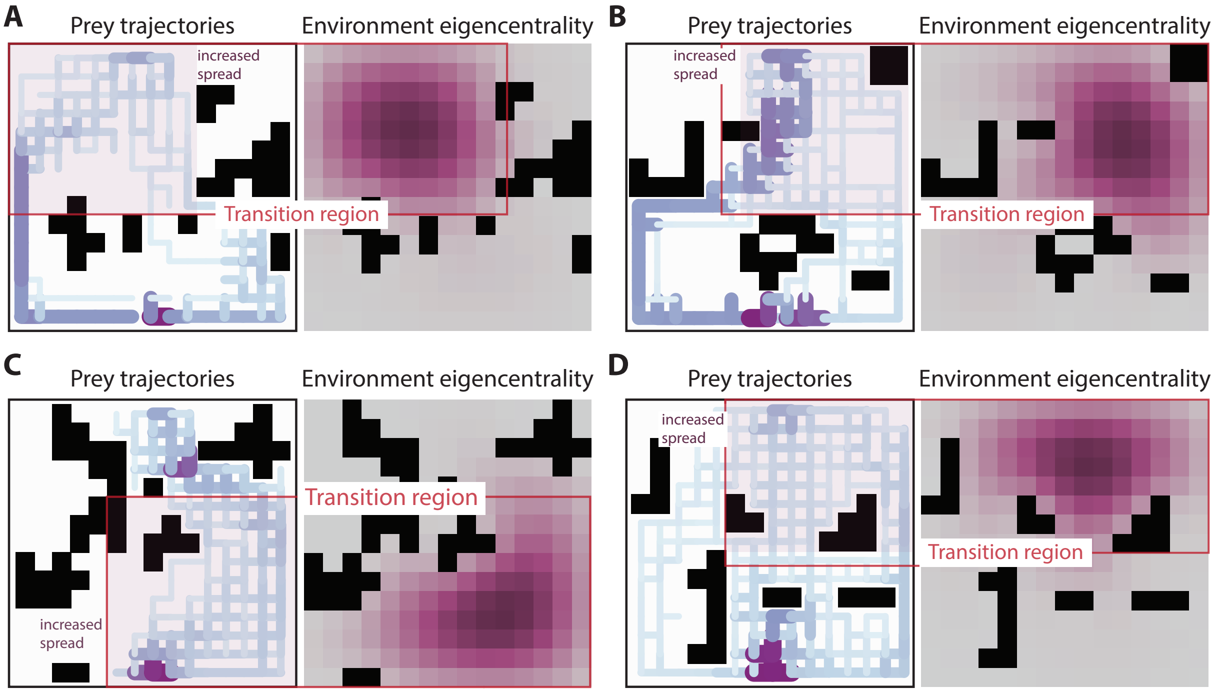


Figure 5: **Example prey success trajectories and corresponding environment eigencentrality.** Example environments and their eigencentralities and eigencentrality gradients. Color densities of each cell are proportional to the metric. Transition regions from low to high eigencentrality based on a change in gradient and value of eigencentrality are shown by the red box. The pink box highlights the increase in path spread that corresponds to regions of high eigencentrality.

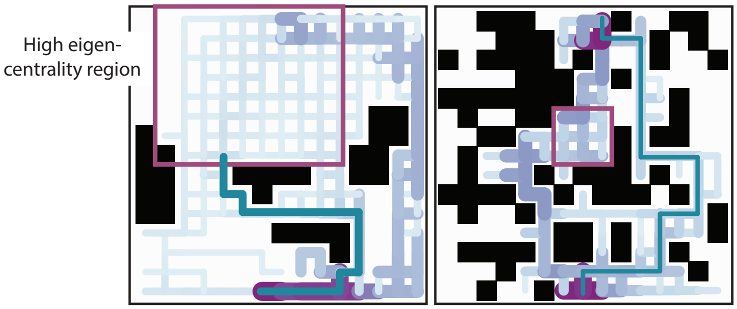


Figure 6: **Example prey success trajectories and corresponding environment eigencentrality.** Heatmaps of all action sequences taken by the prey that resulted in prey survival at the maximum planning level (5000 states), with color density proportional to frequency. Color bar action frequencies range from 1 for low, to 92 for the most saturated color, dependent on entropy level. The pink box indicates the high eigencentrality region.

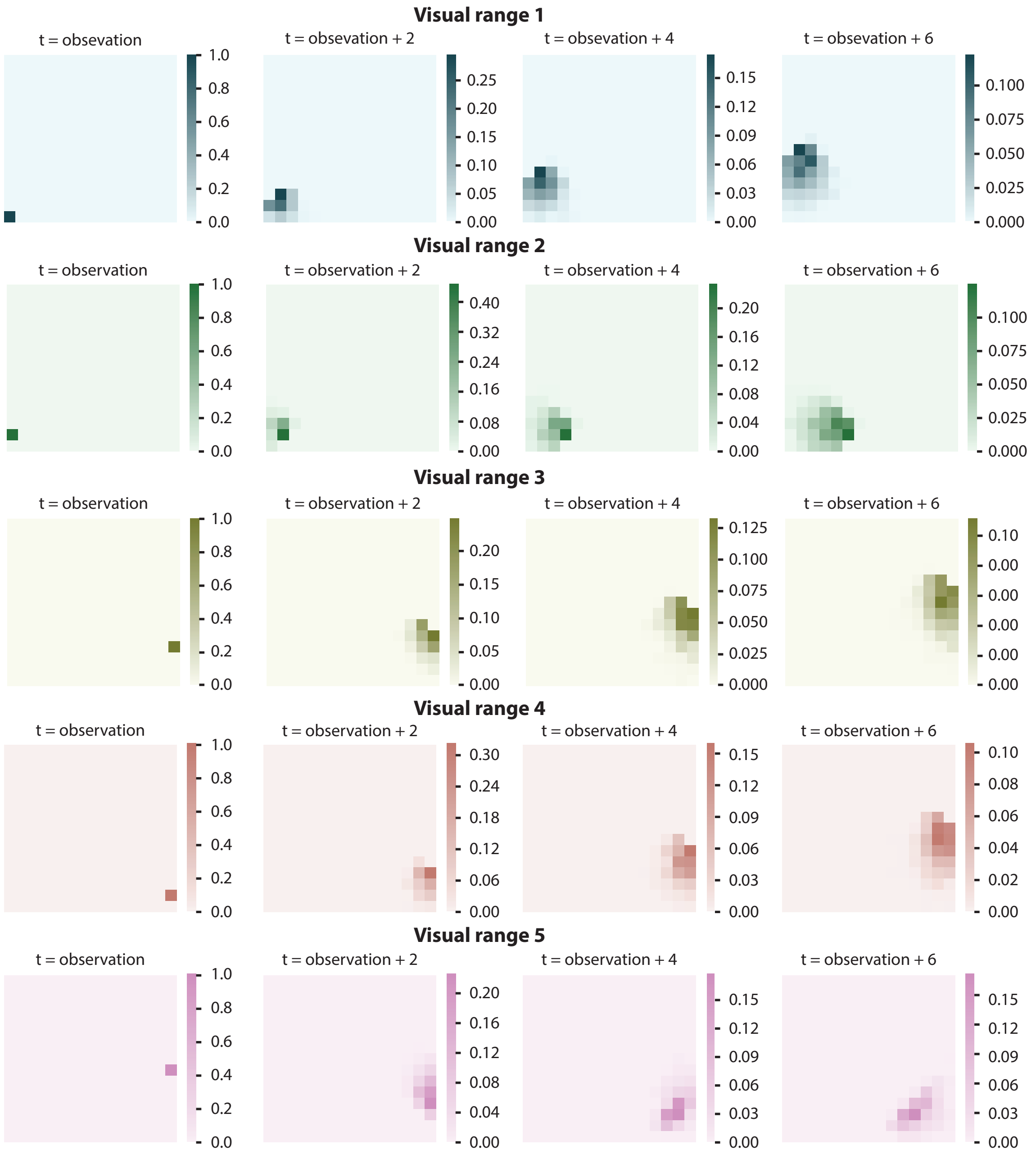


Figure 7: **Uncertainty growth as a function of visual range.** The heatmaps depict the spread of prey's belief state based on its knowledge about how the predator moves (both speed and randomness). The variance of the prey's belief state changes with the number of time steps that the predator is out of the prey's visual cone, and the prey's visual range. A random episode with at least 6 unobserved time steps was chosen, and the policy implemented by the prey from the onset of observation to 6 time steps ahead was implemented. During the prey's implementation of this action sequence, based on the initial predator observation, the predator location was randomly sampled and moved according to the predator model ($N = 1000$).

### 4 Supplementary Tables

Table 1: General parameters used for all simulations

| Parameter | Value | Note |
|---|---|---|
| Environment size | 15 × 15 | The environment size was chosen to have both a large state-space and be feasible in terms of duration of simulations given our compute time on the cluster used for this study (Methods). While the advent of terrestriality increased visual range by a factor of 100, within this environment we were only able to simulate an increase of at most a factor of 5 due to this size limitation for computational expediency. The $x$- and $y$-coordinate values for location are $\in [0, 14]$, with the origin at the bottom left. |
| Prey initial location | (7, 0) | Prey initial location was chosen arbitrarily and was fixed across all computational experiments |
| Predator initial location | | Predator initial location was chosen randomly from unoccupied cells. |
| Safety location | (14, 7) | Safety location was chosen arbitrarily and was fixed across all computational experiments |
| Predator pursuit strategy | 75% aggressive, 25% random | Pursuit probabilities were chosen arbitrarily, however the pac-man example provided in Silver et al.'s work (*5*) also uses this same pursuit strategy. The randomness places importance on the prey observing the predator. After each observation the randomness in predator pursuit strategy causes belief state diffusion (**Supplementary Fig. 7**). In the case of 100% aggressive pursuit, after one observation the prey would know exactly where the predator is after each subsequent time step. |

Continued on next page

Table 1 – continued from previous page

| Parameter | Value | Note |
|---|---|---|
| Predator:Prey speed ratio | 1.5 | With 50% probability (drawn from a Bernoulli distribution) the predator moves 2 cells ahead. Therefore, on average the predator is 1.5× faster than the prey. This speed falls within typical terrestrial predator pursuit speeds relative to prey of 1.2–2 (*1, 2*). |
| Rewards | -1, -25, -100, +1000 | At each time step the prey is required to move; movement of one square received a reward of -1. If the prey hit the wall or an occlusion it received a reward of -25. If the predator occupied the same cell at the same time step as the prey the episode terminates and the prey receives a reward of -100 ("death"). If the prey is able to reach the safety position the episode terminates and the prey receives a reward of +1000 ("survival"). These values are from Silver et al. (*5*). |
| Tested environment entropies | | Entropy 0.0: 0±0 occlusions ; Entropy 0.1: 3±0 ; Entropy 0.2: 9±1 ; Entropy 0.3: 13±1 ; Entropy 0.4: 20±2 Entropy 0.5: 28±3; Entropy 0.6: 35±3 ; Entropy 0.7: 45±3 ; Entropy 0.8: 59±4 ; Entropy 0.9: 75±3 |
| Number of states the prey forward simulates | | Fixed within trial, but varying across trials, this parameter took these values: 1, 10, 100, 1000, 5000. |

Table 2: Parameters for POMCP (1.1.2 **Simulation 1: Algorithm of plan-based action selection**)

| Parameter | Symbol | Value | Note |
|---|---|---|---|
| Exploration constant for PO-UCT | $c$ | 100 | During the within tree search (over nodes that have been visited) the value of an action is augmented by an exploration bonus that is highest for rarely tried actions $\tilde{Q}(s,a) = Q(s,a) + c\sqrt{\frac{\log N(s)}{N(s,a)}}$. This scalar constant determines the relative ratio of exploration to exploitation. Thus, when $c = 0$ the explored nodes are selected greedily. The value chosen was set to the value used by Silver et al. (*5*). Changing the value of $c$, especially lowering it to close to $0$, could possibly change the survival rates, but is unlikely to change the overall trend we see. |
| Softmax inverse temperature parameter | $\beta$ | 3 | Experimental work on human and animal decision making has found that the stochasticity in the observed behavior can be captured by softmax policies (*7, 47*). The parameter $\beta$ controls how diffuse the probability of choosing an action is. As $\beta \to \infty$ the actions are chosen greedily, and at $\beta = 0$ actions are chosen randomly. This seemingly low $\beta$ value is a consequence of the sizes of the rewards: very negative at death, and very positive at survival. This value ensures that most action are chosen greedily from the constructed tree. Increasing $\beta$ would cause more actions to be chosen greedily, and though unlikely, could increase survival rate, but similar to before, would not change the overall observed trend, since the same $\beta$ value is used in all simulations. |

Continued on next page

Table 2 – continued from previous page

| Parameter | Symbol | Value | Note |
|---|---|---|---|
| Discount factor | $\gamma$ | 0.95 | The discount factor affects the weight of the future rewards. Therefore, $\gamma = 0$ will result in state or state-action values to represent only the immediate reward. On the other hand, $\gamma = 1$ will make the agent consider the cumulative effects of long term rewards. Lowering the discount value will decrease the agents cumulative reward at survival, and will increase the impact of the negative reward given at each time step. Thus, this might cause behaviors such as the agent going towards the predator, since the overall imagined cumulative reward might end up being lower. Increasing the $\gamma$ won't have any foreseeable effect, although $\gamma = 1$ could have divergence problems. The value used was taken from the experiments done by Silver et al. (*5*). |
| Rollout count | | $200 \times 1.25^{\text{Env. entropy}*10}$ | In POMCP if a node has not been previously visited in order to determine the value of the node, random action rollouts are carried out. The rollout, similar to the actual simulation, terminates if the prey reaches the safety position or if the predator catches the prey. Therefore, the only stipulation is for the rollout count to be high enough so that the prey encounters either of the termination conditions. In environments where there is not a lot of clutter around 200 was found to be more than enough. When clutter is added to the environment, the nature of random action rollout, causes the agent to need more time steps. The above modulation to the base line of 200 random rollouts was found to be sufficient. |

Table 3: Parameters for PRQL (1.1.3 **Simulation 1: Habit-based action selection**)

| **Parameter** | **Symbol** | **Value** | **Note** |
|---|---|---|---|
| Softmax temperature parameter | $\tau$ | 0 | This parameter controls how diffuse the probability of choosing a policy from the policy library is. Initially $\tau$ is set to $0$, therefore the first chosen policy from the policy library is always random. In consequent trials $\tau$ is updated by: $\tau = \tau + \Delta\tau$. This value was set to the value used by Fernandez et al. (*8*). |
| Incremental increase of $\tau$ | $\Delta\tau$ | 0.001 | This parameter controls how fast the softmax temperature parameter increases. As $\tau \to \infty$ policies from the policy library are chosen greedily based on their weight. Given reward values, this increment allows for continual exploration after one implementation. If this value was increased, it would inhibit the agent from exploring other policies which might also result in favorable outcomes. Decreasing this value would result in policies to be chosen randomly over more trials. This value was set to the value used by Fernandez et al. (*8*). |
| Discount factor | $\gamma$ | 0.95 | Was kept the same as the discount factor that was used in plan-based action selection (**Table 2**). |

### 5 Supplementary Movies

In all videos, the prey is teal, the predator is yellow, the goal location is red, if present visual range is pink, and if present the occlusions are black.

Movie 1: **Representative episodes showing prey behavior with varying visual ranges.** For each visual range (except visual range 1) a representative survival and death trial are shown. The typical form of successful strategies observed independent of visual range can be characterized as wall-following behavior, or thigmotaxis. At high visual ranges, the prey can quickly correct initially incorrect actions it took (going towards the predator). These trials are representative examples, for full survival paths seen with prey with varying visual ranges see **Supplementary Figure 2**.

Movie 2: **Representative low entropy (0.0, 0.1, 0.2, 0.3) episode showing prey wall-following behavior.** For each entropy a representative survival and death trial are shown. The typical form of successful strategies observed in low entropy environments, which can be characterized as wall-following behavior, or thigmotaxis. These trials are representative examples for each environment, for a representation that includes all others see **Supplementary Figure 3** (entropy = 0.0–0.3).

Movie 3: **Representative high entropy (0.7, 0.8, 0.9) episode showing the constriction of successful strategies.** For each entropy a representative survival and death trial and shown. The prey's survival strategy is dependent on the possible escape routes to safety, which are few in number due to the high level of clutter. The trajectories within high entropy environments are correspondingly highly stereotyped. For typical success paths of other high entropy environments, see **Supplementary Figure 3** (entropy = 0.7–0.9).

Movie 4: **Representative mid-entropy strategies.** Different survival strategies (straight towards the goal, hiding, roundabout around occlusions), and an example of a death trial are shown. In mid-entropy environments we observe a diversity of prey strategies even when the predator start location is kept constant. These videos in an environment with entropy = 0.4. Notably, these episodes depict flexible behaviors that are generated to strategically use occlusions based on the current predator strategy.

Movie 5: **Representative mid-entropy strategies.** As in Movie 4, but now in a different environment with entropy 0.5 (occupancy frequency map for this environment across all trials shown in **Supplementary Figure 3**, row 5 of column 6). These videos show different strategies that the prey takes (roundabout around occlusions, broken-wing, hiding).

Movie 6: **Representative mid entropy episode showing different behavioral strategies.** As in Movie 4, in a different environment with entropy 0.5. These videos show the different strategies (hiding, following low eigencentrality, straight towards the goal) that the prey takes.

Movie 7: **Representative mid-entropy episode showing different behavioral strategies with respect to different predator start locations.** These videos show how the predator start location influences prey strategy (full occupancy frequency map is shown in **Fig. 3E3**). For each predator start location an example death episode is always also shown.
